# Phylogenomics supported by geometric morphometrics reveals delimitation of sexual species within the polyploid apomictic *Ranunculus auricomus* complex (Ranunculaceae)

**DOI:** 10.1101/2020.01.07.896902

**Authors:** Kevin Karbstein, Salvatore Tomasello, Ladislav Hodac, Franz G. Dunkel, Mareike Daubert, Elvira Hörandl

**Affiliations:** University of Goettingen, Albrecht-von-Haller Institute for Plant Sciences, Department of Systematics, Biodiversity and Evolution of Plants (with Herbarium), Untere Karspuele 2, D-37073, Goettingen, Germany; University of Goettingen, Georg-August University School of Science (GAUSS), Wilhelmsplatz 1, D-37073, Goettingen, Germany; Am Saupurzel 1, D-97753 Karlstadt, Germany

**Keywords:** Europe, geometric morphometrics, RADseq, *Ranunculus auricomus*, sexual species, target enrichment

## Abstract

Species are the basic units of biodiversity and evolution. Nowadays, they are widely considered as ancestor-descendant lineages. Their definition remains a persistent challenge for taxonomists due to lineage evolutionary role and circumscription, i.e., persistence in time and space, ecological niche or a shared phenotype of a lineage. Recognizing and delimiting species is particularly methodically challenging in fast-evolving, evolutionary young species complexes often characterized by low genetic divergence, hybrid origin, introgression and incomplete lineage sorting (ILS). *Ranunculus auricomus* is a large Eurasian apomictic polyploid complex that probably has arisen from the hybridization of a few sexual progenitor species. However, even delimitation and relationships of diploid sexual progenitors have been unclearly ranging from two to twelve species. Here, we present an innovative workflow combining phylogenomic methods based on 86,782 parameter-optimized RADseq loci and target enrichment of 663 nuclear genes together with geometric morphometrics to delimit sexual species in this evolutionary young complex (< 1 Mya). For the first time, we revealed a fully resolved and well-supported maximum likelihood (ML) tree phylogeny congruent to neighbor-net network and STRUCTURE results based on RADseq data. In a few clades, we found evidence of discordant patterns indicated by quartet sampling (QS) and reticulation events in the neighbor-net network probably caused by introgression and ILS. Together with coalescent-based species delimitation approaches based on target enrichment data, we found five main genetic lineages, with an allopatric distribution in Central and Southern Europe. A concatenated geometric morphometric data set including basal and stem leaves, as well as receptacles, revealed the same five main clusters. We accept those five morphologically differentiated, geographically isolated, genetic main lineages as species: *R. cassubicifolius* s.l. (incl. *R. carpaticola*), *R. flabellifolius*, *R. envalirensis* s.l. (incl. *R. cebennensis)*, *R. marsicus* and *R. notabilis* s.l. (incl. *R. austroslovenicus*, *R. calapius*, *R. mediocompositus, R. peracris* and *R. subcarniolicus*). Our comprehensive workflow combing phylogenomic methods supported by geometric morphometrics proved to be successful in delimiting closely related sexual taxa and applying an evolutionary species concept, which is also transferable to other evolutionarily young species complexes.

## Introduction

Species are the basic units of biodiversity and evolution, but their definition remains a persistent challenge for taxonomists (Sukumaran & Knowles, 2017). Most modern authors consider species as ancestor-descendant lineages (De Queiroz, 2007; Sukumaran & Knowles, 2017), a concept that is applicable to both sexual and asexual lineages. Nevertheless, its rather challenging to define their evolutionary role and circumscription, i.e. persistence in time and space, ecological niche or a shared phenotype (Freudenstein & al., 2017; Hörandl, 2018). Hörandl (2018) proposed a species delimitation workflow based on four main principles particularly addressing apomictic polyploid complexes: First, separate and classify the obligate sexual progenitor species; second, merge sexual progenitors and highly facultative apomictic lineages into one species; third, classify the main clusters of the remaining facultative apomicts as species; and fourth, treat obligate apomictic lineages as agamospecies.

However, recognizing and delimiting even the sexual progenitor species is methodically challenging in fast-evolving, evolutionary young species complexes (e.g. Burgess & al., 2015). Commonly, they are characterized by low genetic divergence, occasionally by hybrid origin, persistent gene flow, introgression and incomplete lineage sorting (ILS). During speciation processes, ILS is rather the rule than an exception causing incongruences in phylogenetic reconstructions (see e.g., Oliver, 2013; Pease & al., 2018). The progenitor species can also be sexual polyploids. Natural hybridization and polyploidy are important evolutionary processes and are regarded as key factors for diversification of Angiosperms (Soltis & Soltis, 2000; Jiao & al., 2011; Alix & al., 2017). Reticulate evolution (e.g., hybridization, allopolyploidy and ILS), however, results in the phenomenon that gene trees do not match species trees (e.g., McBreen & Lockhart, 2006; Pirie & al., 2009).

The application of molecular markers has helped to reconstruct not only relationships of sexual progenitor species but also hybrid origin, reticulate relationships and population genetic structure of apomictic lineages within species complexes (Dobeš & al., 2004; Hörandl, 2004; Paun & al., 2006a, 2006b; Kirschner & al., 2015). Nevertheless, most complexes are evolutionarily young and many originated or diversified in the context of Pleistocene glaciations. Therefore, genetic divergence among taxa is usually quite low. Traditional genetic markers (for example internal transcribed spacer (ITS) markers) often failed to reconstruct fully resolved phylogenies (e.g., Hörandl & al., 2009; Krak & al., 2013; Kirschner & al., 2015). Population genetic markers like microsatellites or AFLPs have been widely used, but are often not homologous among more divergent taxa (Freeland & al., 2011). Hence, species-level relationships in apomictic complexes often remained unresolved.

The application of phylogenomic methods provides a magnitude more loci compared to traditional genetic markers and can resolve relationships of taxa with less than few million (or even less than 100.000) years of divergence (Pellino & al., 2013; Tripp & al., 2017; Tomasello et al., in subm.). Currently, two main approaches are used in species-level systematics: Target enrichment and restriction site-associated DNA sequencing (RADseq; and the similar Genotyping-by-Sequencing = GBS method). Target enrichment (i.e., nuclear genes selected out of a reference genome) is most useful for diverged species and genus-wide relationships (Folk & al., 2015; Schmickl & al., 2016; Tomasello, 2018). RADseq has repeatedly proven its efficiency in resolving intraspecific relationships, phylogenies of closely related species, but also within genera (relationships between less than 100.000 and more than 50 Mya years; (Baird & al., 2008; Hipp & al., 2014; Cavender-Bares & al., 2015; Tripp & al., 2017; Wagner & al., 2018; Pätzold & al., 2019). Although RADseq yields much more loci and SNPs than target enrichment, it is considered to become less informative with increasing species divergence due to increasing mutations in enzyme cutting sites leading to loss of loci (Rubin & al., 2012; Eaton & al., 2017). Specific pipelines were developed for processing (filtering and assembling) of RADseq data, e.g. the IPYRAD pipeline that accounts for indels and is therefore suited for phylogenetic analysis (Eaton, 2014). Several parameters can be optimized to find homologous loci and to maximize the true phylogenetic signal leading to fully-resolved topologies (Paris & al., 2017; McCartney-Melstad & al., 2019; Pätzold & al., 2019). Since incongruences due to reticulate evolution are hard to determine in concatenated alignments, new methods have been developed to examine the phylogenetic tree topologies for conflicted and/or poorly informed branches (e.g. Quartet Sampling (QS) method; Pease & al., 2018).

With a target enrichment approach, single copy loci are selected beforehand based on their position in the genome and to harbor a certain amount of variation. Loci obtained by target enrichment are usually longer compared to those from RADseq allowing for gene trees estimation and allele phasing. The importance of the latter has been highlighted in recent studies (Eriksson & al., 2018; Andermann & al., 2019). They demonstrate the advantages derived by retrieving allele information for a correct phylogenetic inference, particularly in recently diverged species complexes where reticulate evolutionary processes play a tremendously important role. The lack of the need for concatenation makes the employment of coalescent-based methods for species tree and species delimitation inference possible. These methods are particularly useful because they are able to accommodate for all the stochastic processes responsible for incongruence among gene genealogies or between genealogies and species phylogenies (e.g., ILS, gene loss and duplication) while reconstructing the correct species tree and species delimitation scenario. Preferable are model-based methods (Rannala, 2015), which are able to simultaneously estimate gene trees and species tree taking into account gene tree uncertainty and different models of sequence evolution (e.g., BEAST (Heled & Drummond, 2010), BEST (Liu, 2008)). On the other hand, model-based methods apply computationally intensive algorithms, and their application on target enrichment dataset (i.e., on hundreds of loci) is still challenging. Several species tree inference methods have been described in the last decade using gene trees as input, and are therefore easily applied in phylogenomic studies (e.g., PhyloNet (Than & al., 2008), STEM (Kubatko & al., 2009), MRL (Nguyen & al., 2012), ASTRID (Vachaspati & Warnow, 2015), ASTRAL (Mirarab & al., 2014)).

The multi-species coalescent theory (Yang & Rannala, 2014) is also beneficial because it offers a framework to probabilistically estimate species boundaries and therefore delimit species in taxonomically challenging complexes. Different methods capable to infer species delimitation under the coalescent have been proposed in the last years (Pons & al., 2006; Leliaert & al., 2009; Zhang & al., 2013; Yang & Rannala, 2014; Jones & al., 2015). Although for more than a decade they have been applied to animal, fungi and algal species complexes, only in the last years they have been increasingly employed on plant systems (Hu & al., 2015; Ruiz-Sanchez, 2015; Toprak & al., 2016; Naciri & al., 2017; Wagner & al., 2017; Hassanpour & al., 2018; Tomasello, 2018).

Phenotypic differentiation is still regarded as an important criterion for species delimitation (e.g., Stuessy, 2009; Freudenstein & al., 2017). In flowering plants, the shape of organs (e.g., of leaves) is an important criterion for species-level classification, and many quantitative approaches are available to discriminate species (e.g., Jensen & al., 2002). In the past, traditional morphological classification as a purely descriptive approach has led to the subjective descriptions of hundreds to thousands of morphotypes as species due to minor morphological differences, especially in apomictic polyploid complexes (e.g., Stace, 1998). Relying on single, ‘diagnostic’ characters, in general bears the danger of erroneous classifications (Stuessy, 2009). Another challenge is the assessment of the phenotypic variation of characters, which is in plants usually very high due to phenotypic plasticity in response to environmental factors (Stuessy, 2009; Gratani, 2014). Geometric morphometrics makes progress in the exact, objective and fine-scale assessment of differences in leaf shape via landmark methods and appropriate multivariate statistics (Hodač & al., 2014, 2018). Geometric morphometrics can support final taxonomic decisions based on NGS data (Kilian & al., 2015) particularly in complexes where morphological differences are hard to assess with traditional morphological classification. So far, there have been few studies applying both approaches to disentangle phylogenetic relationships in plants (e.g., Jiang & al., 2019).

*Ranunculus auricomus* is one of the largest European apomictic polyploid complexes and a model system to study apomixis and the evolution of phylogenetically young groups (Hörandl, 1998; Hörandl & al., 2009; Hodač & al., 2014; Barke & al., 2018). The distribution ranges from Greenland, Europe, Western Siberia to Alaska, and from the arctic, temperate to the Mediterranean zone (Jalas & Suominen, 1989; GBIF Secretariat 2017). In Europe, species inhabit a broad range of habitats - from stream- and riversides, forests to marshy, humid and semi-dry anthropogenic meadows. The complex comprises more than 800 species (currently 840 taxa in Euro+Med database; Hörandl & Raab-Straube, 2015), mainly described based on a morphological species concept. Hybridization and polyploidization strongly influenced the evolutionary history of *R. auricomus* species: The huge number of morphological diverse polyploid (mainly tetraploid) agamospecies probably arose from hybridization of the few sexual progenitor species (Paun & al., 2006b; Hörandl & al., 2009; Hodač & al., 2014, 2018; Hojsgaard & al., 2014). Already Linnaeus (1753) recognized the existence of two contrasting morphotypes. He described them as *R. auricomus* L. characterized by deeply divided basal leaves from Western Europe and *R. cassubicus* L. characterized by undivided basal leaves from

Siberia (Kvist, 1987). Afterward, several authors realized that these core morphotypes include numerous variants and are interconnected by several hybridogenic intermediates. Because of too many intermediates, a concept of ‘main species’ and agamic lineages as subspecies (Marklund, 1961, 1965) failed. Later on, these main species or morphogroups were no longer formally accepted (Ericsson, 1992; Hörandl, 1998). Phylogenetically, these morphogroups are not monophyletic (Hörandl & al., 2005, 2009). Most of the species (about 600) were described during the 20^th^ century from Northern Europe (Marklund, 1961, 1965; Julin, 1980; Ericsson, 1992). In recent years, some new regional and local species have been recognized in Central Europe (e.g., (Soó, 1964, 1965; Borchers-Kolb, 1985; Hörandl & Gutermann, 1998c, 1999; Dunkel, 2015; Dunkel & al., 2018).

Despite the huge species diversity of apomictic taxa (> 800), only twelve sexual species have been so far described. *Ranunculus cassubicifolius* Koch (1939) is a widespread but disjunctly distributed diploid species (with locally distributed sexual autotetraploid populations) occurring along stream-/riversides, swampy or humid forests and meadows in the Northern and Southeastern edge of the Alps (Hörandl & Gutermann, 1998b; Hörandl & Greilhuber, 2002). *Ranunculus carpaticola* Soó (1965) inhabits humid beech, hornbeam and oak forests and has a disjunct distribution in the Carpathians (Paun & al., 2006a, 2006b; Hörandl & al., 2009). The two species form tetra- and hexaploid, apomictic hybrids lineages distributed in C to NE Slovakia (Hörandl & al., 2009). *Ranunculus flabellifolius* Rchb. (1832) is a conspicuous species probably restricted to the Western Romanian Carpathians (Jalas & Suominen, 1989; Dunkel & al., 2018). Nearby, a few occurrences were also mentioned outside of the Carpathians in Serbia (Stevanović & al., 1991). Similar to *R. carpaticola* and *R. cassubicifolius*, *R. flabellifolius* is characterized by non-dissected leaves but differs from them in fan-shaped leaves (due to connate segments), and the infrequent occurrence of up to three-lobed basal leaves.

The other sexual species share dissected basal leaves and a heterophyllous basal leaf cycle. *Ranunculus marsicus* Guss. & Ten. (1835), *R. envalirensis* Grau (1984) and *R. cebennensis* Dunkel (2018) are high mountain species with small, restricted geographical ranges. Diploid *R. envalirensis* inhabits alpine meadows in the Eastern Pyrenees, *R. cebennensis* is only known from two different meadows in Massif Central (France), and the only (mainly) tetraploid sexual species *R. marsicus* (also with penta- to heptaploid asexual cytotypes) occurs in the Central Apennines in alpine meadows. The other dissected basal leaf species are restricted to the Illyrian region, a warm humid deciduous forest area ranging from Southeastern Austria to Central and Southwestern Serbia (see e.g., Košir & al., 2008). The diploid *R. notabilis* Hörandl and Guterm. (1998) was first described from Southeastern Austria and is characterized by strongly dissected leaves. *Ranunculus calapius, R. austroslovenicus*, *R. mediocompositus*, *R. peracris* and *R. subcarniolicus* grow in Slovenia and Western Croatia, partly in close sympatry, and were recently described (Dunkel & al., 2018). Obligate sexual reproduction was ascertained previously for *R. marsicus* (tetraploids) by Masci & al., (1994) based on embryological studies, for *R. cassubicifolius*, *R. carpaticola* and *R. notabilis* using embryology and flow cytometric seed screening (FCSS) by Hojsgaard et al. (2014, and previous literature therein) and Barke et al. (2018), and for the other taxa included here by Dunkel et al. (2018) using FCSS and/or pollen quality measurements. For other European species of the complex, apomixis was confirmed using embryological (Häfliger, 1943; Nogler, 1984) or cytological data (Rousi, 1956), large-scale FCSS screenings on more than 200 populations (Karbstein et al., in prep.), and pollen quality measurements (Hörandl & al., 1997; Hörandl & Gutermann, 1998a, 1998c, 1998b, 1999; Dunkel, 2010, 2011).

Applying isozyme markers, the split between sexual species of both contrasting morphotypes *(R. carpaticola*/*cassubicifolius* and *R. notabilis*) was estimated 900,000 years ago whereas the split between morphological similar *R. carpaticola* and *R. cassubicifolius* approximately occurred 300,000 years ago (Hörandl, 2004). The *R. auricomus* group as a whole is monophyletic within the genus and related to North American and Central Asian taxa (Emadzade & al., 2011, 2015). Molecular phylogenetic investigations based on genomic data covering all diploid and tetraploid sexual species of the complex have not been conducted, and hence information on the monophyly and the degree of genetic divergence of taxa is lacking. Thus, the delimitation of sexual species within this complex is taxonomically unclear.

Species within the *R. auricomus* complex exhibit various basal leaf shapes throughout the year, but only the strongest dissected spring basal leaf is the taxonomically the most important one (Borchers-Kolb, 1985; Hörandl & Gutermann, 1995, 1998a). Furthermore, only the central part of the lowermost stem leaf is deemed to be of taxonomic relevance (Hörandl & Gutermann, 1995). Geometric morphometrics has been shown to elucidate morphological leaf diversity of experimental F_2_ hybrids between the morphologically most divergent species (Hodač & al., 2018). However, a concatenated analysis using all taxonomically informative traits of all sexual species has not yet been conducted.

Consequently, we address the following questions in our study: Is a combination of RADseq and target enrichment able to resolve distinct evolutionary lineages in the sexual taxa of the *R. auricomus* complex? Do these lineages match described species, or do we have to revise the species delimitation? Do phylogenetic relationships match similarity patterns found by geometric morphometrics? Can we propose a well-founded species concept based on NGS data supported by geometric morphometrics? Finally, we aim to establish an innovative workflow combining phylogenomics (RADseq and target enrichment) and geometric morphometrics to delimit species in evolutionary young complexes using the *R. auricomus* group as a model system.

## Materials and Methods

### Study locations and population sampling

We included all twelve described sexual species of the *Ranunculus auricomus* complex in the present study (Table 1). From 1998 to 2018 (mainly 2017 and 2018), we sampled up to 20 individuals per population and one to three populations per species in Europe including locations in Andorra, Austria, Croatia, France, Germany, Hungary, Italy, Romania, Slovakia to Slovenia (see Table 1, Fig. 1). We used the diploid species *R. pygmaeus* and *R. sceleratus* as outgroup species according to the *Ranunculus* phylogeny of Emadzade et al. (2011). We sampled living plants and kept them in the Old Botanical Garden at the University of Goettingen under controlled environmental conditions for further flow cytometric, genetic and geometric morphometric analyses. Herbarium specimens were also collected for all populations and deposited in GOET. Additionally, we recorded altitude, GPS coordinates and habitat characteristics of populations. We dried fresh leaves in silica gel for further flow cytometric and genetic laboratory work. Individuals were cultivated in 1.5 l pots with Fruhstorfer Topferde LD 80 (garden beds without differences in solar radiation and water supply).

**Fig. 1.**
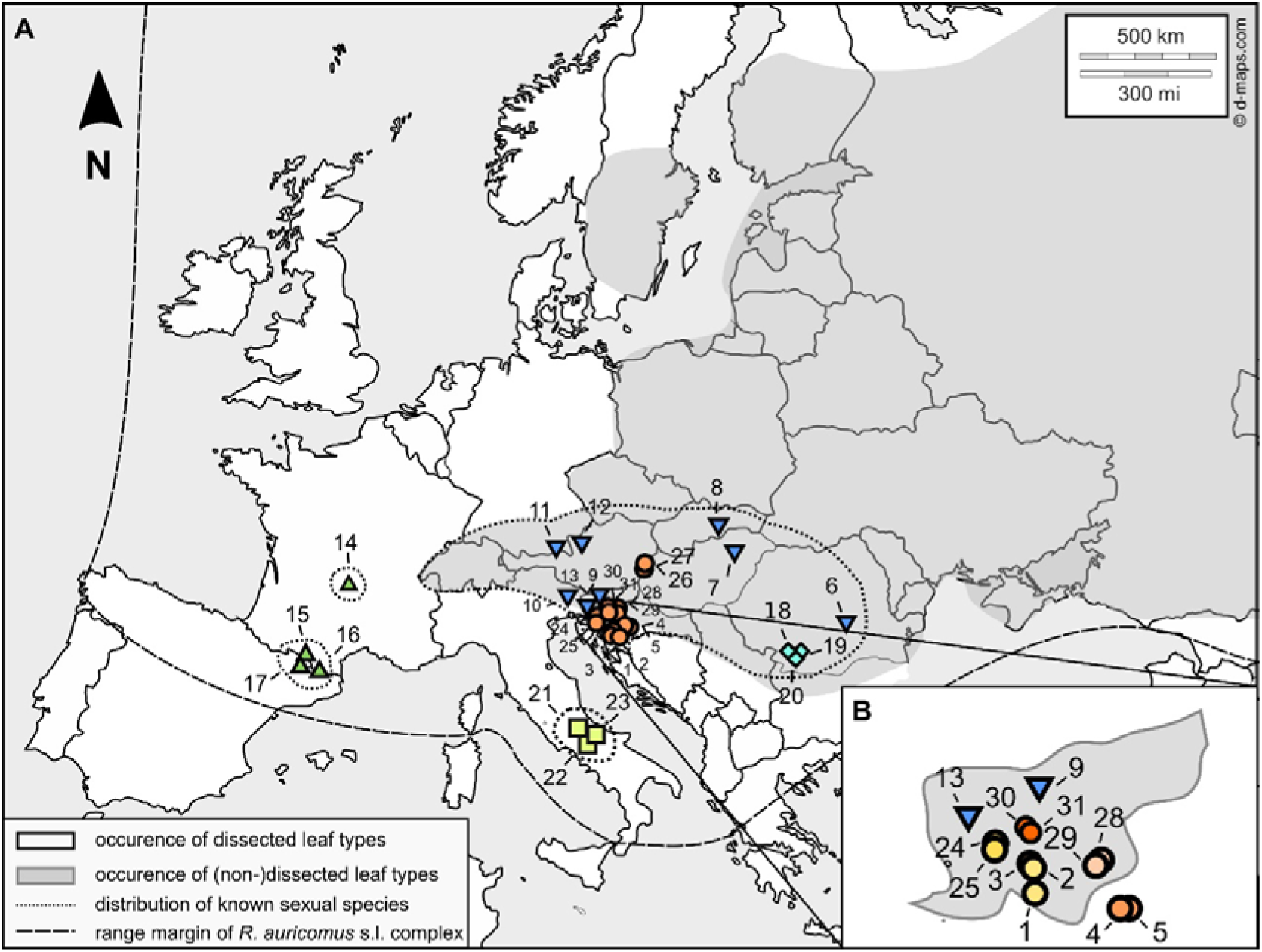
(A) Study locations of the 28 sexual *R. auricomus* populations across Europe. The range margin of the whole *R. auricomus* complex is indicated by a broad dashed line whereas the distribution of the known sexual species is marked with a narrow dotted line. (B) Study locations of the sexual *R. auricomus* populations in the Illyrian region (Slovenia, Croatia). Shades of color represent the occurrence of non-dissected (grey) and/or dissected (white) leaf morphotypes (Jalas & Suominen, 1989; GBIF Secretariat 2017). Locations are indicated by circles with different numbers corresponding to locations in Table 1. Colors of circles represent populations belonging to one accepted species. We downloaded the original map from https://dmaps.com/m/europa/europemax/europemax-10.svg.

**Table 1.**
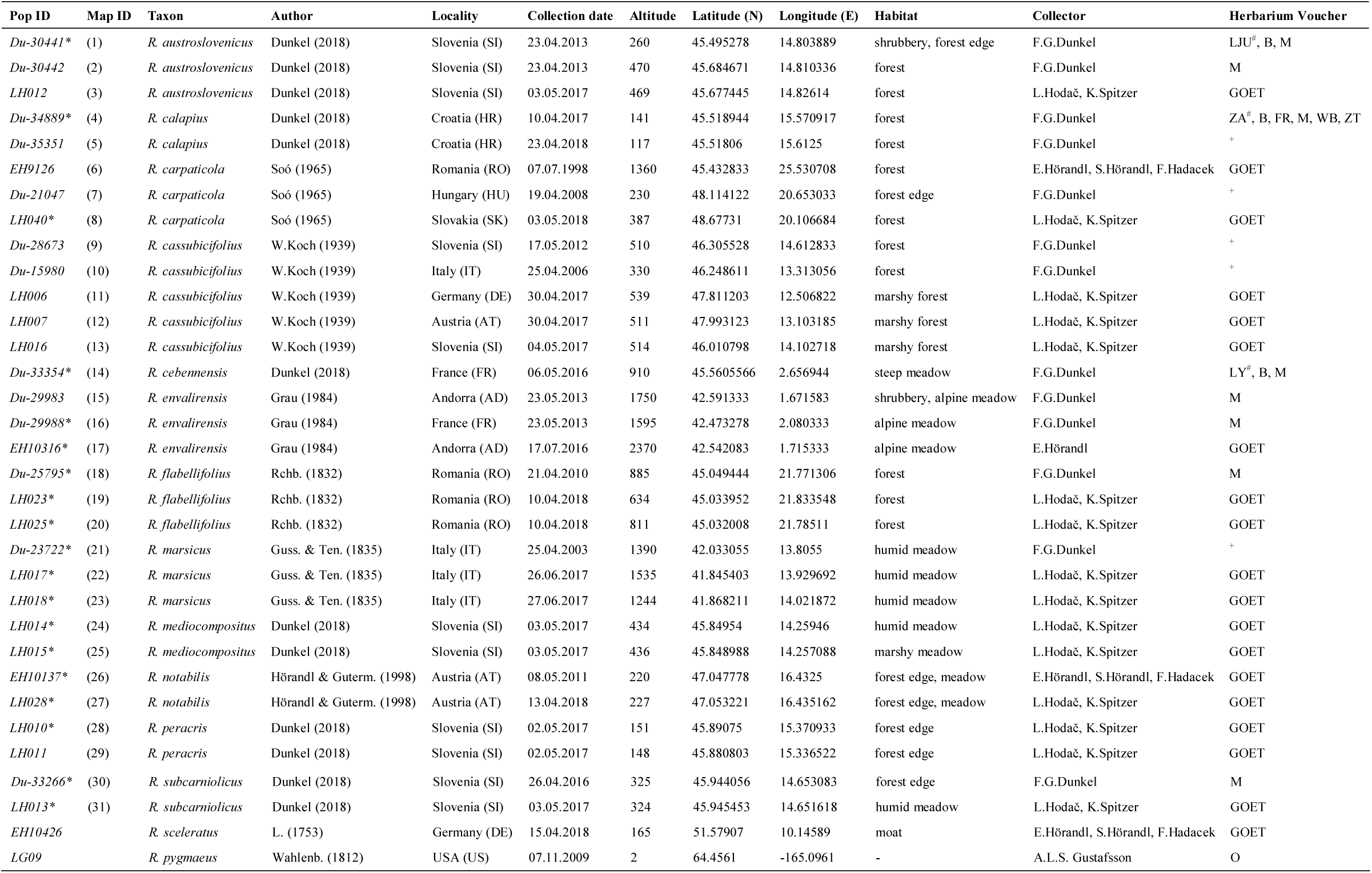
Locations of 31 studied populations of sexual diploid and tetraploid species within the *R. auricomus* complex across Europe. Population ID with map ID (see Fig. 1), taxon name, name author, locality by country and ISO code 3166-2, collection date of population sampling, altitude (in meter above sea level, m.a.s.l.), latitude (N, decimal), longitude (E, decimal), habitat, collector and herbarium voucher specimens. F. G. Dunkel (Karlstadt, Germany) kindly provided garden individuals and habitat characteristics of populations with sample number ‘Du’. * locality is at locus classicus or nearby, ^#^ holotype, ^+^ private herbarium F.G.Dunkel.

### Flow cytometry (FC) and single seed flow cytometric seed screening (ssFCSS)

We checked ploidy levels of all samples conducting flow cytometry (FC) of DNA material dried in silica gel and collected freshly in spring 2017 to 2019. Silica dried leaf material (∼0.5-1 cm²) was ground to small pieces by one steel ball (Ø 4 mm) in a 2 ml Eppendorf tube with Tissue Lyzer II (Qiagen, Hilden, Germany; 30 Hz s^−1^, time 7-14 s) basically following Klatt et al. (2016) and Barke et al. (2018) (see Supplementary Table 1). Analyses were carried out for one to 15 individuals per population yielding 191 FC measurements in total (see FCSS Table in the data repository). Data were quality filtered according to the following settings: A maximum coefficient of variation of 6.0, a minimum number of parts within a particular peak of 500 and a minimum number of parts in total measurement of 4000. We removed all samples that did not match the quality settings.

For confirmation of the sexual mode of reproduction, we conducted single seed flow cytometric seed screening (ssFCSS) using seeds harvested in summer 2017 to 2019. We analyzed seeds as described in FC measurements but with slight modifications: We added 100 µl extraction buffer Otto I to the seed and ground the seed for 5-7 s. Then, we put 100 µl extraction buffer Otto I to the ground seed material and immediately inverted samples for 30 s. Afterward, we filtered samples through CellTrics® filters with a mesh of 30 µm into flow cytometric sample tubes and stained them with 800 µl Otto II. We conducted ssFCSS analysis for five to 10 seeds per individual and one to three individuals per population yielding five to 23 seeds per population (see FCSS Table in the data repository). In total, we performed 205 FCSS measurements. We set a maximum coefficient of variation of 7.0, a minimum number of parts within a particular peak to 100 (embryo) and 300 (endosperm), and a minimum number of parts in total measurement of 3000 as quality thresholds.

All analyses were performed on a CyFlow Ploidy Analyzer (Sysmex, Norderstedt, Germany). We used the diploid *Ranunculus auricomus* individual ‘J6xF7/01’ (Barke et al. 2018) as an external standard and for gain adjustment, i.e., the voltage of the photomultiplier (Doležel & al., 2007). The software ‘CUBE16’ vers. 1.6 (Sysmex, Norderstedt, Germany) was used to calculate the DNA content (as relative fluorescence intensity) by determining the mean peak ratios of standard and leaf sample, coefficient of variation (CV) and number of parts (per peak, and in total). To ascertain the reproductive pathway, we calculated the ratio between endosperm ploidy and embryo ploidy as peak index (PI). A PI around 1.5 indicates a sexual pathway because a diploid embryo and a triploid endosperm formed after double fertilization. Peak indices of 2 or higher indicate apomictic pathways, as the unreduced egg cell develops parthenogenetically while the endosperm developed without fertilization or with contribution of one or two reduced sperm nuclei (PI = 2, 2.5 and 3.0, respectively; Matzk & al., 2000; Doležel & al., 2007; Klatt & al., 2016). If the embryo was still in development and hence the peak too small for a reliable FCSS measurement, we used the leaf value of the mother plant to calculate the ratios (see FCSS Table in the data repository).

We carried out statistical analysis in R vers. 3.6.1 using the graphical user interface RSTUDIO vers. 1.1.383 (R Foundation for Statistical Computing, 2019). We calculated mean leaf/embryo peak values for each taxon.

### Laboratory work - DNA extraction, RADseq and target enrichment

We extracted the total genomic DNA of 45 *R. auricomus* samples from ∼1.5 cm² silica-dried leaf material using the Qiagen DNeasy Plant Mini Kit® (Qiagen, Hilden, Germany). Additionally, we also extracted silica-dried leaf material of the outgroup species *Ranunculus sceleratus* and *R. pygmaeus*. We followed the manufacturer’s instructions but prolonged sample incubation in lysis buffer to one hour increasing DNA yield. We checked DNA concentration using the Qubit® fluorometer and the Qubit® dsDNA BR Assay Kit (ThermoFisher Scientific, Waltham, USA). We diluted DNA solutions to 30 ng/μl in a target volume of 55 μl, checked DNA quality by gel electrophoresis, and removed samples with low quality (fragmented bands). The samples packed in dry ice were sent to Floragenex Inc. (Portland, USA) for generating RAD libraries based on the protocol of Baird & al. (2008). The enzyme *PstI* was used for digestion, and the multiplexed samples were sequenced on an Illumina® HiSeq 4000 platform to produce 100 bp single-end reads at the University of Oregon Genomics Core Facility. We checked the quality of raw reads using FastQC vers. 0.11.8 (Andrews, 2010).

For target enrichment, we used almost the same samples as in RADseq analyses (additional tetraploid *R. cassubicifolius* samples, see Table 1; Tomasello & al., subm.). Probe design for target enrichment was done from RNA sequences of *R. carpaticola*, *R. cassubicifolius* and *R. notabilis* published by Pellino et al. (2013) and is described in detail in Tomasello et al. (subm.). In total, the baits system consisted of 17,988 probes (14,632 of which unique), used to in-liquido capture 736 target genomic regions. Library preparation and hybrid capture protocol are described in detail in the supplements (Supplementary Text 1). Two different paired-end sequencing runs (2x 250 bp) were performed on an Illumina MiSeq System (Illumina Inc., San Diego, USA). The obtained raw reads were processed using the pipeline HybPhyloMaker (all scripts available at: https://github.com/tomas-fer/HybPhyloMaker/; Fér and Schmickl (2018). Before analyses, reads were phased using SAMtools vers. 0.1.19 (Li & al., 2009) in order to retrieve allelic information. After processing data on HybPhyloMaker and filtering for missing data (see Supplementary Text 1), we were able to retrieve information from 663 of the initial 736 nuclear loci (2495 exons in total) excluding 73 loci (see Supplementary Text 1). Then, these alignments were used for the subsequent gene tree/species tree and coalescent-based species delimitation analyses. Although chloroplast sequences were generated during laboratory work, we excluded them because sequences were not informative to resolve species relationships.

### *De novo* assembly of RAD loci and parameter optimization in IPYRAD

The following analyses were performed with IPYRAD vers. 0.7.28 (Eaton, 2014) on the GWDG HPC Cluster (Goettingen, Germany). Raw reads were demultiplexed allowing no mismatch in barcode sequence. Afterward, raw reads were cleaned by removing adapter sequences and restriction overhang (TGCAG) and trimmed if the Phred quality score was below 20 (maximum low-quality base calls in a read). Reads shorter than 35 bp were excluded after adapter trimming. For parameter optimization, we developed a workflow accounting for different ploidy levels of sexual *R. auricomus* species. First, we separated diploid and tetraploid species into two datasets; second, we (separately) optimized in-sample clustering threshold (ISCT, within individuals) threshold of diploid and tetraploid species; third, we optimized between-sample clustering (BSCT, between individuals) threshold of merged diploid and tetraploid assemblies; and fourth, we selected the optimal minimum number of samples per locus by choosing the maximum likelihood tree with best bootstrap support. We included all 45 individuals in the optimization process, and basically followed strategies of parameter optimization described in Paris et al. (2017) and Pätzold et al. (2019) optimizing ISCT and BSCT separately. For each dataset (2n, 4n), we kept all parameters constant (default) except minimum depth for statistical/majority rule base calling and clustering threshold (minimal sequence similarity in percent) for *de novo* assembly. We specified the minimum depth to 6 and 12, and the range of clustering threshold from 60 % to 99 % (60, 70, 80, 85, 90, 91, 92,…, 99).

To optimize ISCT, we carried out calculations for different minimum depths and clustering thresholds and analyzed the output regarding the number of clusters within samples, average cluster depth and clusters rejected due to high heterozygosity (Supplementary Fig. 1A–F). With decreasing ISCT threshold, no. of clusters (steepest between 0.99 - 0.96 (2n) / 0.99 - 0.93 (4n)) and cluster depth (steepest between 0.99 - 0.96 (2n) / 0.99 - 0.93 (4n)) decreased, and clusters rejected due to high heterozygosity increased (steepest between 0.94 - 0.60 (2n) / 0.91 - 0.60 (4n)) (Supplementary Fig. 1). We selected an ISC threshold of 0.95 for diploid species and an ISC threshold of 0.92 for tetraploid species minimizing rejected clusters and maximizing no. of clusters and cluster depth (see Pätzold et al. (2019). Generally, fewer clusters of mindepth12 compared to mindepth6 setting were rejected due to high heterozygosity. Therefore, we selected the mindepth12 setting. Then, we merged the optimized diploid and tetraploid assemblies. To optimize BSCT, we again conducted calculations for different minimum depths and clustering thresholds (setting maximum alleles to 2) and analyzed the output regarding number of polymorphic loci, SNPs, loci filtered by maxSNPs (maximum of allowed SNPs), removed duplicates (two ISC loci merged into one BSC locus), shared loci and new polymorphic loci (Supplementary Fig. 1 G–L). Then, we selected the optimal BSCT value. With decreasing ISCT threshold, no. of polymorphic loci, SNPs and shared loci steeply increased (peak ∼ 0.95 - 0.90) followed by a smooth decrease (Supplementary Fig. 1). No. of loci filtered by maxSNPs and removed duplicates raised in a sigmoidal manner with decreasing BSC threshold (turning point ∼ 0.93 - 0.95). From BSCT 0.99 to 0.95, the merged assembly gained loci, from 0.94 to 0.90 no. of polymorphic loci pended around 45000 followed by a smooth decrease. We chose a BSC of 0.92 for the merged assembly minimizing no. of removed duplicates and loci filtered by maxSNPs and maximizing no. of polymorphic loci, SNPs and shared loci (see also Paris & al., 2017 and Pätzold et al. (2019). Generally, fewer loci were filtered by maxSNPs and removed duplicates of mindepth12 compared to mindepth6 setting. Therefore, we selected the mindepth12 setting. Furthermore, we assessed the impact of different minimum no. of samples per locus on missing data, no. of loci (resulting alignment), and maximum likelihood tree bootstrap values (Supplementary Fig. 2). With decreasing minimum no. of samples per locus, amount of missing data (3.7 - 85.3 %), no. of loci (134 - 205738) and average bootstrap support (until ‘min10’ setting) increased remarkably. No of loci drastically ascends but average bootstrap support slightly decreased from ‘min10’ to ‘min4’ setting. Moreover, the topology did not differ from ‘min50’ setting downwards. Therefore, we picked out the ‘min10’ dataset as the optimized assembly (see also workflow Fig. 2).

To evaluate the impact of missing loci on the topology of phylogenetic inferences, we performed the final filtering step in IPYRAD specifying the minimum number of samples to 2 (4%), 5 (10%), 14 (30%), 24 (50%), 33 (70%), and 42 (90%). We assessed the number of loci and the number of missing data for each step (Supplementary Fig. 2).

**Fig. 2.**
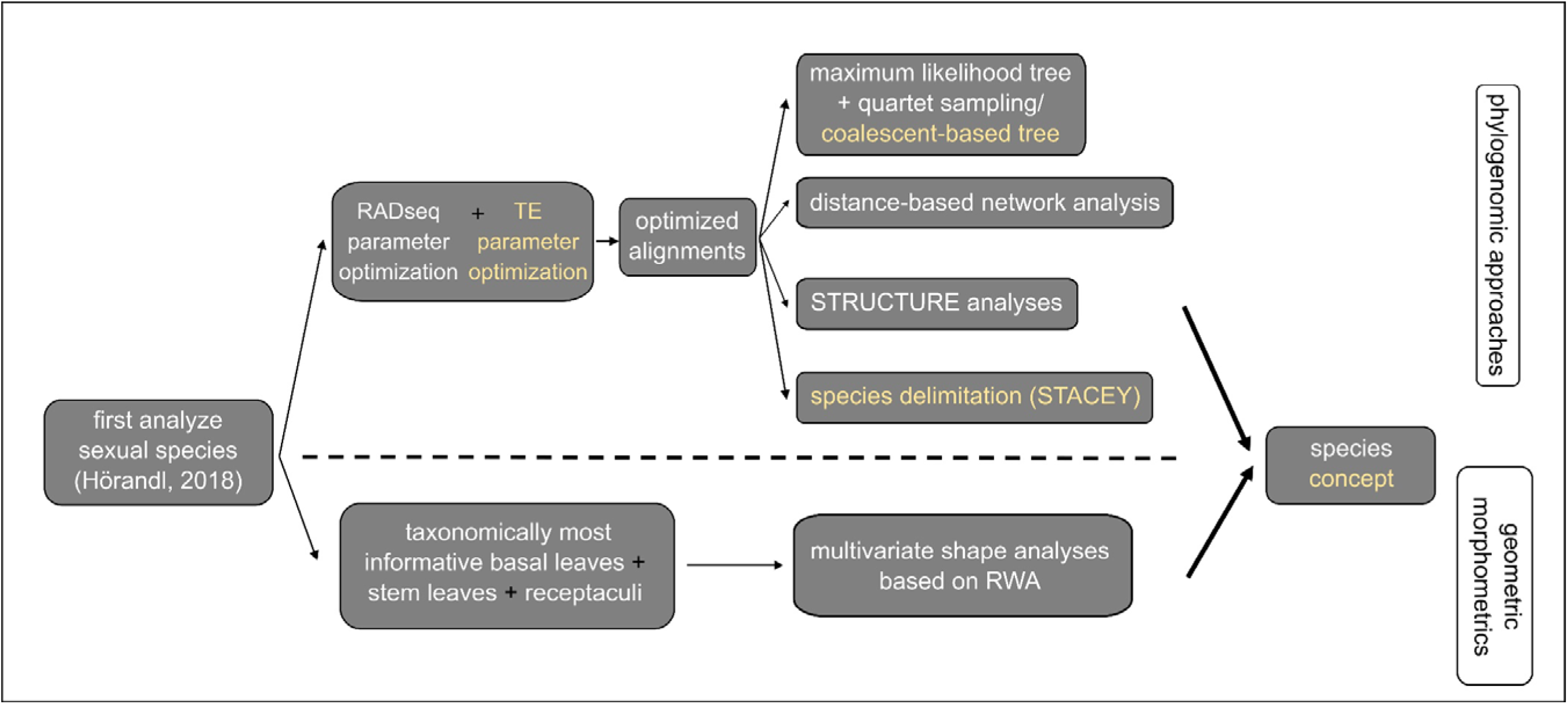
The basic workflow of this study: As proposed by (Hörandl, 2018) for polyploid apomictic species complexes, we first analyzed the sexual species of *R. auricomus*. We carried out a dual-approach based on phylogenomic data supported by geometric morphometrics to propose a well-defined species concept. Phylogenetic inference was mainly based on an optimized RADseq dataset (see Methods for details): We conducted a ML analysis and investigated node support with the quartet sampling (QS) method (Pease & al., 2018). Structure and distance-based network analysis (Neighbor-net) examined the clustering/grouping of individuals. We used an optimized target enrichment (TE) data set to calculate a coalescent-based species tree and a species delimitation analysis (STACEY; Bouckaert & al., 2014). We included the taxonomically most informative traits (basal leaves, stem leaves and receptacles) to carry out multivariate shape analysis based on RWA.

### Phylogenomic analyses - Maximum likelihood tree and quartet sampling of RAD loci

Maximum likelihood (ML) analyses were performed on all resulting alignments (4%, 10%, … and 90%) with EXAML vers. 3.0.21 (Kozlov & al., 2015; Fig. 2). Here, we only used *R. sceleratus* as an outgroup because extractions from *R. pygmaeus* from herbarium material did not reveal sufficient high-quality DNA for RADSeq. We used the new rapid hill-climbing algorithm and set the gamma model of rate heterogeneity and a random number seed. Additionally, we enabled the memory saving option for gappy alignments regarding data sets that contained more than 50% missing data (4 - 30 % min. loci per sample; see Supplementary Fig. 2). We also ran RAXML vers. 8.2.4 (Stamatakis, 2014) to generate 100 bootstrap replicate alignments with their respective parsimony starting trees. Subsequently, we performed maximum likelihood analyses with EXAML on every replicate alignment based on the above-specified options. Then, we plotted the results on the EXAML tree (full dataset). For the following analyses, we selected the alignment ‘min10.phy’ (minimum number of samples: 5 (10%); Supplementary Fig. 2) due to the highest mean bootstrap support among all ML trees. We visualized all resulting trees in FIGTREE vers. 1.4.3 (Rambaut, 2014).

To examine the phylogenetic tree for conflicted and/or poorly informed branches, we conducted the Quartet Sampling (QS) method (Pease & al., 2018). We set 100 replicates per branch, number of threads to 4 and log-likelihood threshold cutoff to 2. Quartet Sampling (QS) method outputs three different node scores (Pease & al., 2018): The quartet concordance (QC) describes the ratio of the concordant to both discordant quartets (1 = all concordant, > 0 = more discordant, 0 > more concordant); the quartet differential (QD) defines skewness of both discordant patterns (1 = equal, 0.3 = skewed, 0 = all topology 1 or 2); and quartet informativeness (QI) characterizes the proportion of informative replicates (1 = all informative, 0 = none informative). QD values around 1 can indicate incomplete lineage sorting (ILS) due to the presence of both discordant topologies (random pattern) whereas values towards 0 potentially hint at directional introgression events due to the presence of one particular alternative topology (Pease & al., 2018). We drew results together with bootstrap values on the ‘min10.phy’ maximum-likelihood tree (Fig. 3). The calculation procedure also gives the quartet fidelity score (QF, see Supplementary Table 2 for details) indicating whether a taxon is misplaced within the topology (1 = not misplaced, 0 = misplaced; see rogue taxon approaches; Pease & al., 2018).

**Fig. 3.**
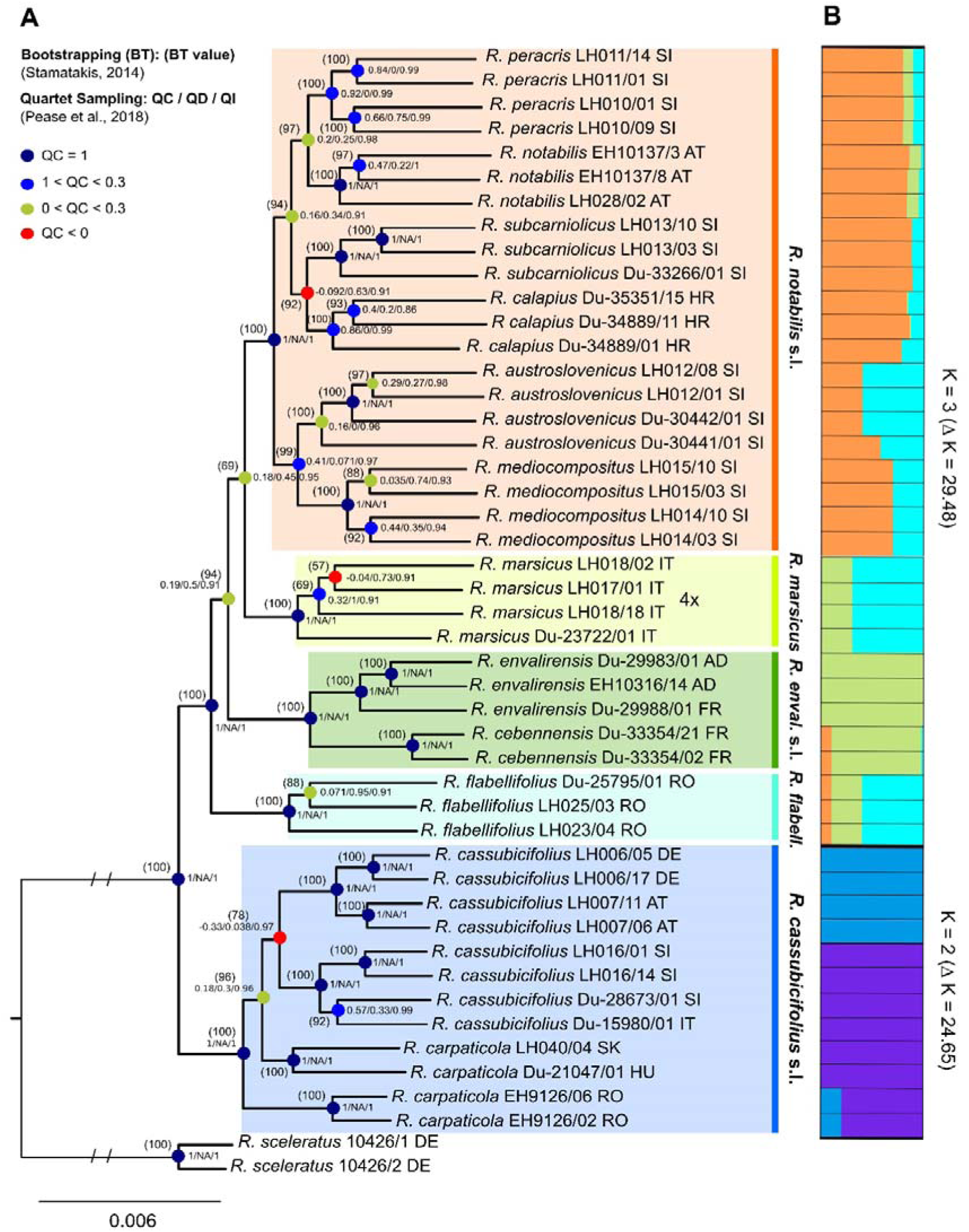
(A) Maximum likelihood (ML) tree of 45 sexual species of the *R. auricomus* complex and *R. sceleratus* as outgroup. Bootstrap values (BT) are given in brackets above nodes. Quartet Sampling scores QC (quartet concordance), QD (quartet differential) and QI (quartet informativeness) are arranged to the right of the node (QC/QD/QI). If QC equals 0, then QD is NA (skewness of discordant patterns cannot be determined if there are no discordant patterns). See Supplementary Table 2 for QF (quartet fidelity) score. Nodes are colored according to QD value (see figure legend). Boxes around clades specify the accepted species with their names. Country labels behind taxa are in ISO code 3166-2 format. (B) Right to ML tree, the bar graphs of the genetic structure analysis for each of the two subsets with the most likely K values are illustrated (see Supplementary Table 3 for results of the Evanno tests).

### Phylogenomic analyses - Genetic distance

In order to test for reticulate (non-tree like) evolution of sexual *R. auricomus* species, we calculated a network analysis based on the neighbor-net algorithm using the program SPLITSTREE vers. 4.14.6 (Huson & Bryant, 2006). To estimate the genetic distance, we specified a general time reversible (GTR) model with estimated site frequencies and maximum likelihood (equal rates of site variation, default rate matrix). Then, we performed bootstrapping with 1000 replicates and showed values > 80 for the main clusters.

### Phylogenomic analyses - Genetic structure

To infer the genetic composition of sexual *R. auricomus* species (Fig. 2), we took the SNP based format including one randomly selected variable site per locus from IPYRAD (‘min10.ustr’; dataset without outgroup). We performed a genetic structure analysis in STRUCTURE vers. 2.3.4 (Pritchard & al., 2000), and set an admixture model, a burn-in of 5000 and a MCMC of 50000 replicates. K was set from one to 11, and each K was replicated five- to 10-times. To determine the optimal K value, we ran the Evanno method implemented in STRUCTURE HARVESTER (Earl & vonHoldt, 2012; see Supplementary Table 3). We used CLUMPP vers. 1.1.2 (Jakobsson & Rosenberg, 2007) to merge all runs of the optimal K value, and plotted genetic structure with DISTRUCT vers. 1.1 (Rosenberg, 2004). Since we observed the deepest split between *R. carpaticola*/*cassubicifolius* and the other species, we conducted two additional IPYRAD analyses (same settings as ‘min10.phy’) considering the above-mentioned groups (subsets). K was set from one to six (*R. carpaticola*/*cassubicifolius*) / to 11 (*flabellifolius* to *peracris*), and each K was replicated five- to 10-times. The choice of K, merging of runs and illustration of results followed the procedure described above (see Supplementary Table 3). The second most likely K values of the total data set and both subsets were usually characterized by a very low delta K value, and were therefore not drawn. We illustrated the results of both subsets together with the ML tree (Fig. 3).

### Phylogenomic analyses - Target enrichment and coalescent-based species delimitation

In addition to the concatenated tree obtained with the RADseq data, we estimated the species tree using the target enrichment dataset and the coalescent-based method ASTRAL (Mirarab & al., 2014). For the scope, gene trees (exon trees) were first reconstructed with ML as implemented in RAxML vers. 8.2.4 (Stamatakis, 2014). We run analyses with 100 standard bootstrap replicates, partitioned by exon, and applying the GTRGAMMA model to all partitions. RaxML trees were rooted and branches with bootstrap values lower than 50 were collapsed using the script ‘Newick Utilities’ (Junier & Zdobnov, 2010). Finally, the species tree was inferred using the coalescent-based algorithm implemented in ASTRAL-III vers. 5.6.3 (Zhang & al., 2018) with 100 multi-locus bootstrap replicates (Seo, 2008).

We used the BEAST2 package STACEY vers. 1.2.1 (Jones, 2017) to infer species boundaries among the sexual representatives of the *R. auricomus* complex (Fig. 2). STACEY uses a Bayesian approach to infer species delimitation and species phylogeny based on the multispecies coalescent model. It is one of the few model-based programs able to perform this type of analysis. STACEY was demonstrated to outperform the other programs in estimating the correct ultrametric species tree (Andermann & al., 2019), or while inferring the best species delimitation scenario (Tomasello, 2018).

We ran BEAUTI vers. 2.5.2 (Bouckaert & al., 2014) to create an input file for STACEY. We used the 50 most informative loci (with the highest number of parsimony informative sites) from those sequenced applying a target enrichment approach by Tomasello et al. (subm.). The alignments (between 500 and 1000 bp long) consisted of a maximum of 56 OTUs (two alleles from 28 samples, outgroups were excluded from the analyses). During the inference, substitution models, clock models and gene trees were treated as unlinked for all loci. Sequence substitution models were selected for each locus separately using the Bayesian Information Criterion (BIC) in jModeltest vers. 2.1.10 (Darriba & al., 2012). In the Stacey .xml input file, parameters of the substitution models were fixed to those found in jModeltest. The strict clock was enforced for all loci fixing to 1.0 the average clock rate of one random locus while estimating all other clock rates in relation to this locus. We set the ‘Collapse Height’ to 1*10^−4^. This parameter has no biological meaning, and values between 0.000001 and 0.0001 usually produce similar results in similar times (Jones, 2017). The ‘Collapse Weight’ parameter was estimated using a Beta prior with parameters α = 1.0 and β = 1.0, which represent a flat distribution between 0 and 1 (i.e., values close to one indicate higher probability densities for low number of species whereas values close to zero higher probability densities for high number of cluster/species). We followed this approach in order to equally investigate all possible scenarios of species delimitation. Finally, we gave to the bdcGrowthRate prior a log-normal distribution (M = 4.6, S = 1.5), a gamma shape (α = 0.1 and β = 3.0) to the popPriorScale prior, α and for the relativeDeathRate we set a beta prior (α = 1.0, β = 1.0).

The input files were run for 1*10^9^ iterations sampling every 1*10^5^th generation. Two independent runs were performed to check convergence among independent analyses. We checked convergence and ESS (values > 200) in Tracer vers. 1.6 (Rambaut & al., 2018). We combined output files containing the trees sampled in the two independent runs using LogCombiner vers. 2.5.1 (Bouckaert & al., 2014) after discharging 10 % of the analyses as burn-in. The obtained file was processed with the ‘species delimitation analyser’ [speciesDA.jar, Jones & al., (2015); available at: http://www.indriid.com/2014/] using a ‘collapse height’ of 1*10 ^−4^ and setting the similarity cut-off to 0.9. Finally, we produced a similarity matrix using a modified version of the R script provided in (Jones & al., 2015).

### Geometric morphometrics

The geometric morphometric data set comprised predominantly material sampled from living plants completed with leaves from herbaria (Supplementary Table 4, Fig. 2). We selected the most taxonomically informative traits for geometric morphometric analyses, i.e., the most dissected basal leaves in the leaf cycle, the central part of the lowermost stem leaf and the receptacle at fruit stage (Borchers-Kolb, 1985; Hörandl & Gutermann, 1995, 1998a; Hodač & al., 2018). We collected one to seven fresh basal and stem leaves per flowering plant in April to May 2018 and 2019, and scanned them immediately after harvesting in 400 dpi resolution using a Epson Perfection V500 Photo (SEIKO Epson Corp., Suwa, Japan) and CanoScan LiDE 220 (Canon Inc., Ōta, Japan) scanner to avoid potential bias of morphometric analysis due to herbarization (Volkova & al., 2010; Klingenberg, 2015). We harvested receptacles at the fruiting time from June to August 2018 and 2019, and digitized them under a Leica M125 microscope (Leica Microsystems GmbH, Wetzlar, Germany) with 10-15fold magnification. In total, we included 592 basal leaves, 580 stem leaves and 421 receptacles in the following data analyses based on 215, 210 and 170 individuals, respectively (see Supplementary Table 4). We only used individuals that are characterized by all three morphological traits for the concatenated data analysis.

We followed Hodač & al. (2018) for image pre-processing and creation of TPS files. Adjusted č specimens have been imported into TpsDig vers. 1.40 (Rohlf, 2015) to perform digitization of 2D landmarks and semilandmarks (Bookstein, 1997a; MacLeod, 2008). We applied three different landmark approaches to each dataset (see Supplementary Fig. 3 for details). For basal leaves, we adopted the landmark configuration from Hodač & al. (2018) and placed 26 landmarks along the leaf outline according to the leaf venation assuring homology of landmarks among samples. Landmarks, i.e., point locations, have the advantage to objectively assess morphological differences between anatomically homologous positions of different samples (Klingenberg & al., 2012; Gunz & Mitteroecker, 2013; Hodač & al., 2018). For stem leaves, we developed a new application by putting eight landmarks along the outline of the middle leaf segment according to the venation of the central vein. Due to a gap of homologous locations, we filled the gap between landmarks two and 13, and 18 and 27 with 10 semilandmarks roughly situated in corresponding positions, respectively. Semilandmarks (or sliding landmarks) are able to capture length to width ratios (Gunz & Mitteroecker, 2013), which is important to stem leaves of sexual species within the *R. auricomus* complex. The spacing of semilandmarks is optimized by sliding along the leaf shape and after sliding, landmarks and semilandmarks can be treated together for statistical analyses (Perez & al., 2006; Gunz & Mitteroecker, 2013). For receptacles, we also developed a new application by putting eight landmarks along the receptacle outline including locations at the androclinium, intervallum, gynoclinium and central uppermost part. We filled the gap between landmarks four and 10, and 10 and 16 with five semilandmarks roughly situated in corresponding positions, respectively. We also used semilandmarks (or sliding landmarks) capturing length to width ratios.

We applied the thin-plate-spline approach (Bookstein, 1997b) to extract shape variables from our landmark-based data. Because *R. auricomus* shows high phenotypic plasticity with extremely divergent forms, we tested the suitability of the thin-plate-spline approach using the program TpsSmall vers. 1.34 (Rohlf, 2015). The first data standardization procedure comprised the separation of symmetric and asymmetric components of shape variation. The landmark configurations were symmetrized (after Klingenberg & al., 2002) with respect to the bilateral symmetry axis defined by landmarks 1+14 in basal leaf, landmarks 1+15 in stem leaves and landmarks 1+10 in receptacles. The symmetrization followed the approach described in Klingenberg et al. (2002) (for details see the capture of Supplementary Fig. 3).

Symmetrized landmark configurations (each representing a single plant) were analyzed in the program TpsRelw vers. 1.70 (Rohlf, 2015) to search for group clustering within basal/stem leaf and receptaculum morphospaces. The resulting scores of specimens (= mean shapes per plant) on the relative warps then represented input shape variables for further analyses. The intra- and interspecific morphological variability has been analyzed separately for basal/stem leaves and receptacles. Specimens have been plotted in the morphospace captured by the 1^st^ and 2^nd^ relative warp. Scores of specimens on up to four relative warps extracted from each morphological structure have been concatenated and used as input datasets in exploratory and discriminant analyses of species differentiation.

The concatenated morphometric dataset included 11 shape variables (basal leaf - 3, stem leaf - 4, receptaculum - 4), which were available without missing data in 108 plants (23 *R. cassubicifolius* s.l., 23 *R. envalirensis*, 7 *R. flabellifolius*, 3 *R. marsicus*, 52 *R. notabilis* s.l.). To assess morphological differentiation among species, we used the concatenated dataset and performed a non-metric multidimensional scaling (NMDS) analysis with Euclidean distances in the program PAST vers. 2.17c (Hammer & al., 2001). We used the same software to test for morphological differentiation among species with non-parametric multivariate analysis of variance (NP-MANOVA) and 10,000 permutations. Important shape variables responsible for species differentiation were visualized together with morphological clusters as a biplot ordination diagram drawn from canonical variates analysis (CVA) aiming in maximizing the differences among a priori predefined groups (i.e., species).

Reconstructions of shape changes described by relative warps (shape principal components) were modeled with the thin-plate-spline method in the program TpsRelw vers. 1.70 (Rohlf, 2015). We used the same method for computation of species mean shapes averaged over plant individuals in the program MorphoJ vers. 1.06d (Klingenberg, 2011). We finally examined how shapes vary with size (allometry). To assess which shape variables exhibit allometry, we used centroid sizes extracted from original unaligned landmark configurations as an independent variable. Shape variables (only symmetric component) were regressed on the independent variable (centroid size) and the significance of the regression model was computed with 1000 permutations in the program MorphoJ vers. 1.06d (Klingenberg, 2011). In addition to the regression model, we have tested the association between size and particular shape variables and computed Spearman’s rank correlation coefficients (ρ) for centroid sizes and scores on the four most important relative warps. Both, centroid sizes and scores on relative warps were exported from MorphoJ vers. 1.06d (Klingenberg, 2011) and correlations were computed in the program PAST vers. 2.17c (Hammer & al., 2001).

### Indumentum of receptacle

Receptaculum trichome density has been regarded as an important differential character by most taxonomists (e.g., Marklund, 1961, 1965; Julin, 1980; Borchers-Kolb, 1985; Hörandl & Gutermann, 1998a; Dunkel & al., 2018). We assessed trichome density as the number of trichomes in a 0.25 mm² transect. We placed a 0.5 mm transect on the receptaculum using the image manipulation software GIMP 2 vers. 2.10.06 (The GIMP Team, 2018) and counted the number of trichomes within the transect for a total of 219 receptacles. We used R vers. 3.5.3 (R Foundation for Statistical Computing, 2019) to illustrate trichome density per species/accepted species (mean/species) with boxplots and to examine group differences with Kruskal-Wallis rank-sum test followed by pairwise Wilcoxon rank-sum tests with holm correction.

## Results

### Flow cytometry (FC) and FCSS

All species were diploid except for *R. marsicus* exhibiting mainly tetraploid individuals (see Supplementary Table 5). *Ranunculus marsicus* population LH017 comprised tetraploid, pentaploid and hexaploid individuals. FCSS results confirmed the sexual reproductive pathway of all diploid taxa and tetraploid *R. marsicus* (LH018). The *R. marsicus* population LH017 exhibited only one sexual seed from a tetraploid individual and asexual seeds from tetraploid, pentaploid and hexaploid individuals (see Supplementary Table 5). The sexual *R. cassubicifolius* population LH016 exhibited also two asexual seeds formed by only one individual.

### Phylogenomic analyses - Maximum likelihood tree and quartet sampling of RAD loci

We retained 7,237 million raw reads in average per sample (range: 1,413 to 14,405 million raw reads). After filtering steps, analyses yielded 7,235 million reads (range: 1,412 to 14,401 million reads). After optimization of the ISC and BSC thresholds, we finally obtained 24,338 loci per sample on average (range: 8,950 - 38,435 loci), and 86,782 loci and 400,271 SNPs for the total data set.

The ML analysis revealed a fully resolved and well-supported phylogeny of the sexual species within the *R. auricomus* complex (bootstrap values = BT, in brackets, Fig. 3). All described morphospecies are monophyletic. The first (and deepest split) separates the clade containing *R. cassubicifolius* and *R. carpaticola* (non-dissected leaf morphotypes) from the clade containing *R. flabellifolius*, *R. envalirensis*, *R. cebennensis*, *R. marsicus* and the Illyrian species (non-dissected and dissected leaf morphotypes). The first main clade is geographically east-west partitioned in a Carpathian group (*R. carpaticola*), a Southern Alpine - Illyrian group and a Northern Alpine group of *R. cassubicifolius* s.str., the latter with low support (BT = 78). Within the second main clade, surprisingly, the species with mostly non-dissected leaves, *R. flabellifolius* mainly found in the Western Romanian Carpathians, appears on the first, well-supported branch (BT = 100). The next branch (BT = 100) contains the high-altitude species *R. envalirensis* from the Western Pyrenees and *R. cebennensis* from the Massif Central in France. The split between *R. marsicus* and the Illyrian clade is less supported (BT = 69). The only known tetraploid sexual *R. marsicus* from the high-altitudes in the Apennines is thus positioned between the other high-altitude species and the Illyrian ones. The Illyrian species cluster in two well-supported subclades: first, *R. austroslovenicus* and *R. mediocompositus* (BT = 99), and second, *R. calapius*, *R. subcarniolicus*, *R. notabilis* s.str. and *R. peracris* (BT = 94). Within these subclades, bootstrap support is partially lowered (see Fig. 3).

We examined ‘placement’ of the taxa and node support regarding incongruences with the quartet sampling method (QC/QD/QI below bootstrap value; Fig. 3): The taxon-specific quartet fidelity score (QF) ranged between 68 and 93% concordant replicates (mean 82%), i.e., taxa are widely ‘placed’ correctly within the phylogeny (see Supplementary Table 2). All nodes were highly informative indicated by quartet informativeness (QI) values between 91 and 100% informative replicates per node (mean 98%). The quartet concordance (QC) value revealed frequently nodes with high amounts of concordant patterns (> 0.3; 63.7%) but also with particular amounts of discordant patterns (< 0.3; 27.3%, Fig. 3). Within the first main clade (*R. carpaticola/cassubicifolius*), the node splitting *R. cassubicifolius* (Northern Alpine group) and *R. cassubicifolius* (Southern Alpine - Illyrian group) showed more discordant than concordant patterns (QC = −0.33) that are skewed towards one alternative topology indicating directional introgression. Particularly the nodes separating the recently described Illyrian species are moderate to highly conflicted. The nodes splitting *R. austroslovenicus* and *R. mediocompositus*, and *R. notabilis* s.str. and *R. peracris* revealed moderate to low amounts of concordant patterns (QC = 0.41, 0.2) but showed a skew towards one alternative topology potentially also indicating (directional) introgression (QD = 0.071 and 0.25). Moreover, the node between *R. calapius* and *R. subcarniolicus* revealed more discordant than concordant patterns (QC = -0.092) potentially rather caused by ILS (QD = 0.63). Generally, low bootstrap values corresponded to low QC values. In contrast, some nodes with high bootstrap values (90 - 100) were moderately conflicted possessing low QC values.

### Phylogenomic analyses - Genetic distance

The clusters of the Neighbor-net analysis confirmed the topology found in the ML analysis, but also revealed discordant, network-like structures among particular taxa (Fig. 4). We found the deepest split among species between the *R. cassubicifolius*/*carpaticola* cluster and all other species. Within the *R. cassubicifolius*/*carpaticola* group, three clusters can be recognized: (1) *R. carpaticola* from the South Carpathians, (2) *R. carpaticola* (Hungary, Slovakia) plus *R. cassubicifolius* s.str. (Southern Alpine - Illyrian group), and (3) *R. cassubicifolius* s.str. (Northern Alpine group). The last two clusters particularly exhibit reticulation events at the branch basis. Three clusters with a similar degree of divergence and high BT support are located between the most distant groups: *R. flabellifolius* (BT = 100), *R. cebennensis* plus *R. envalirensis (*BT = 100), and *R. marsicus* (BT = 99.8). Interestingly, the Illyrian cluster arises from one basis (BT = 88), and its subclusters show a low genetic distance with strong reticulations on the basis. Subclusters are *R. austroslovenicus*, *R. mediocompositus* and *R. subcarniolicus*, and *R. calapius*, *R. peracris* and *R. notabilis*.

**Fig. 4.**
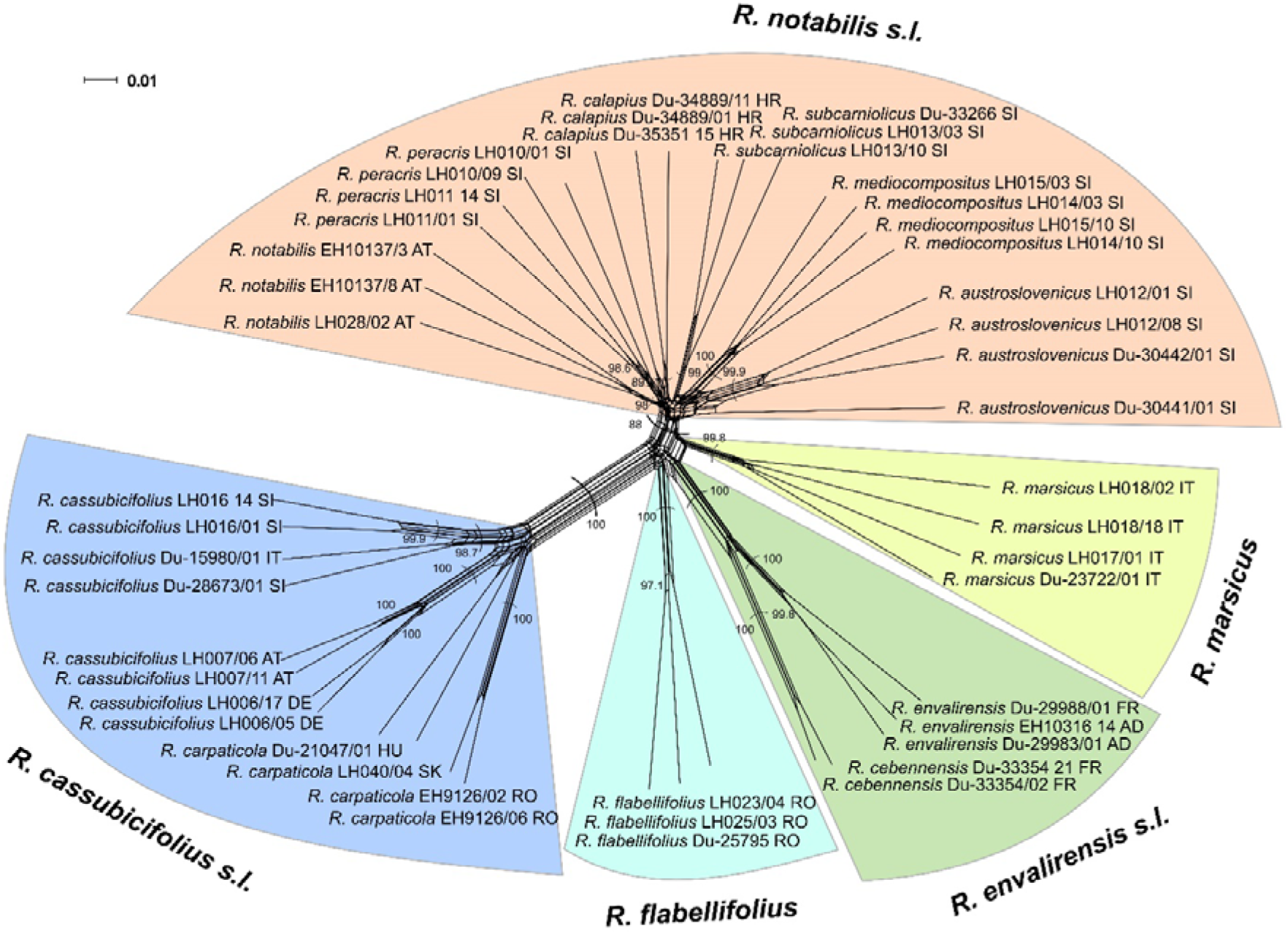
Neighbor-net analysis of 45 sexual samples of the *R. auricomus* complex based on distances of a General Time Reversible (GTR) model with estimated site frequencies and ML (equal rates of sites variation, default rate matrix). We show bootstrap values above 80 for major genetic clusters. We highlighted major clusters with colors and designated them with accepted names. Country labels behind taxa are in ISO code 3166-2 format. Scale bar = no. of changes.

### Phylogenomic analyses - Genetic structure

The likeliest K value of the total dataset (diploids and tetraploids) is K = 2 (ΔK = 2582.67) as ascertained by the Evanno method. The bar graph of the most likely K value exhibit two main genetic partitions (Supplementary Fig. 4): A cluster containing *R. carpaticola* and *R. cassubicifolius* s.str. (corresponding to the first main clade in the ML tree), and a partition containing *R. flabellifolius*, *R. envalirensis*, *R. cebennensis*, *R. marsicus* and the Illyrian species (corresponding to the second main clade in the ML tree). Individual genotypes showed no admixture.

The subset containing *R. carpaticola* and *R. cassubicifolius* s.str. is composed of K = 2 genetic partitions (ΔK = 24.65): *R. carpaticola* (Carpathians) and *R. cassubicifolius* s.str. (Southern Alpine - Illyrian group) form one genetic partition (violet), the Northern Alpine *R. cassubicifolius* the other one. Two *R. carpaticola* individuals from the Southern Carpathians possess also a small fraction from the Northern Alpine group) partition (Fig. 3). The second main clade revealed K = 3 genetic partitions (ΔK = 29.48). Almost all individuals exhibit admixture (Fig. 3). *Ranunculus flabellifolius* is characterized by three genetic partitions, *R. marsicus* by two, and both species have a main genetic partition in common (cyan in Fig. 3). *Ranunculus envalirensis* and *R. cebennensis* share a main genetic partition (green in Fig. 3) although the latter showed a small proportion of admixture coming from the Illyrian group and *R. flabellifolius* or *R. marsicus*. The genotypes of the Illyrian species share a big proportion of the third genetic partition (orange in Fig. 3). The other two genetic partitions of this group appear in smaller, varying proportions.

### Phylogenomic analyses - Target enrichment and coalescent-based species delimitation

Results from the ASTRAL species tree estimation (Fig. 5) are comparable to the tree topology obtained with the RADseq data. The main incongruences between results from the two datasets are (i) the position of *R. marsicus*, and (ii) relationships among described species in the Illyrian clade. *Ranunculus marsicus* is found to be sister to the Illyrian group in the RADseq analyses, whereas the same position is occupied by the *R. cebennensis* - *R. envalirensis* clade in the ASTRAL tree. However, these relationships are characterized by very short, weekly supported branches in both analyses. Relationships among taxa of the Illyrian clade are also contrasting in the two reconstruction methods. Although samples from the same described species are found to be monophyletic (all except for *R. austroslovenicus* in the ASTRAL tree), branches describing relationships among these taxa received low support in both analyses.

**Fig. 5.**
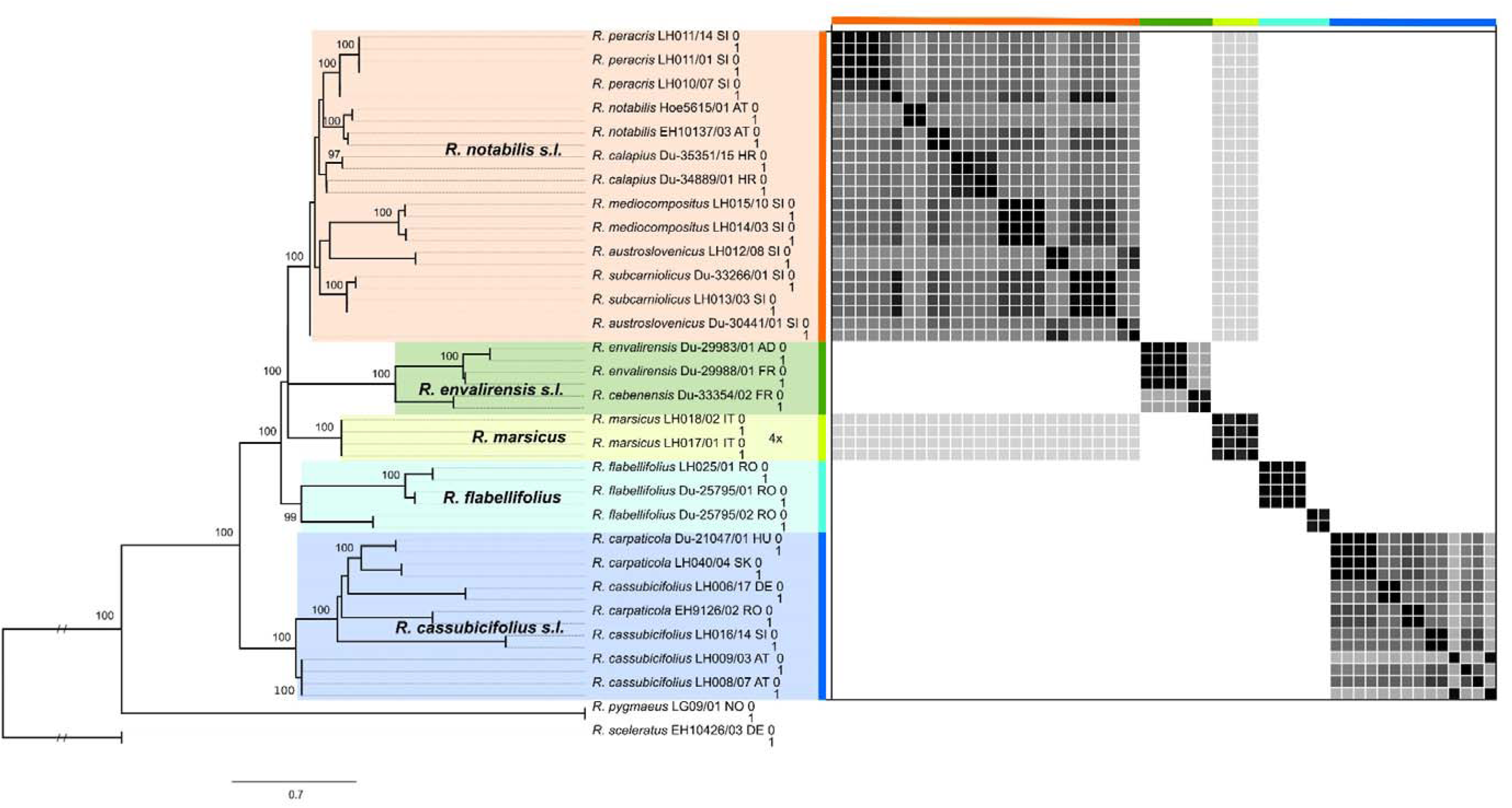
Coalescent-based species tree and species delimitation analyses of sexual species within the *R. auricomus* complex. Species tree from ASTRAL (left) was obtained using values information from 2495 exon trees. Bootstrap values above 95 are shown above branches. In the obtained species tree, each accession was represented by one leaf, and they were duplicated here for graphical purposes. Colored boxes around clades reflect the here proposed taxonomic treatment (as in Fig. 3, the RADseq ML tree). Country labels behind taxa are in ISO code 3166-2 format. On the right part of the figure, there is the similarity matrix obtained from the STACEY analyses. Posterior probabilities to belong to the same cluster (species) are shown for pairs of individuals. Black is for 1.0 posterior probability, white for 0.0.

The STACEY species delimitation analysis preferred scenarios with a reduced number of species. The ‘species delimitation analyser’ found classifications with six genetic clusters (species) as the ones with the highest frequency (0.11). The similarity matrix (Fig. 5) showed six well-defined clusters: Samples of 1) *R. cassubicifolius* and *R. carpaticola,* 2) *R. flabellifolius,* 3) the remnant *R. flabellifolius*, 4) *R. marsicus*, 5) *R. envalirensis* and *R. cebennensis*, and 6) all Illyrian taxa. In almost all cases, individuals from a cluster have zero posterior probability of belonging to another cluster. The only exception is the tetraploid *R. marsicus*, which samples show alternatively a discreetly high posterior probability of belonging to the same cluster of the Illyrian taxa. The cluster containing *R. carpaticola* and *R. cassubicifolius* is weakly differentiated and showed no grouping according to morphospecies or geographical regions. Interestingly, samples from *R. flabellifolius* formed two distinct, well-separated clusters although occurring mainly in a small, restricted area in the Western Romanian Carpathians. The cluster containing the high-altitude species of the Pyrenees and Massif Central exhibit a weak differentiation between *R. envalirensis* and *R. cebennensis*. Even though samples from the Illyrian taxa are not clearly separable, a certain structure is recognizable within their cluster. For example, *R. mediocompositus*, *R. calapius* and *R. peracris* form respectively well-defined subclusters, whereas *R. mediocompositus* shows a remarkable higher similarity to *R. subcarniolicus* than to the other Illyrian taxa. In contrast, samples from *R. notabilis* cluster separately.

### Geometric morphometrics - Morphological gradients

Relative warps analysis of basal leaves (Fig. 6) pointed out a major differentiation among species following a shape gradient from dissected leaves (*R. envalirensis*, *R. marsicus*, *R. notabilis* s.l.) to non-dissected leaves (*R. cassubicifolius* s.l., *R. flabellifolius*). Regarding great variation and overlap in morphospecies, we focus in the following on differentiation of the five main genetic lineages (see above). The best separating warp (bl1) with 56% of the described variance was significantly correlated with leaf size (Spearman’s ρ = −0.6; *P* < 0.001), i.e., divided leaves were smaller and undivided ones were bigger. The second relative warp (bl2) with 18% of the described variance showed a leaf shape change from wide blade bases and narrow lateral segments to narrow blade bases and wide lateral segments. This morphological gradient was not associated with leaf size separating species with divided leaves, namely *R. envalirensis* and *R. notabilis* (Fig. 6A). The third (bl3; 9%) and fourth (bl4; 5%) relative warps described shape changes concerning the middle and first lateral segment (Supplementary Fig. 3A) and altogether the warps bl1-bl4 captured 88% of the variation in basal leaves. The major morphological gradient in stem leaves (sl1) with 72% of captured variation separated *R. cassubicifolius* s.l. from the most other species (Fig. 6B). The shape change from narrow non-dissected to wide dissected stem leaves was positively correlated with size (Spearman’s ρ = 0.2; *P* < 0.001). The second stem leaf warp (sl2; 15%) was not correlated with size. It described shape change from wide non-dissected to narrow dissected stem leaves and separated slightly *R. flabellifolius* from the remaining species. The warps sl1-sl4 explained 97% of the variation in stem leaves. Receptacles exhibited the lowest separating power and neither the first relative warp (rt1; 46%) nor the second (rt2; 34%) could clearly separate the species (Fig. 6C). The first relative warp revealed shape change from smaller wide receptacles to narrow bigger receptacles (Spearman’s ρ = 0.4; *P* < 0.001). In addition, the second warp exhibited allometry (rt2; Spearman’s ρ = 0.6; *P* < 0.001) describing shape change from smaller receptacles with wide and high androclinium and short gynoclinium to bigger receptacles with short and narrow androclinium and high gynoclinium (Fig. 6C). The warps rt1-rt4 explained 92% of the variation in receptacles.

**Fig. 6.**
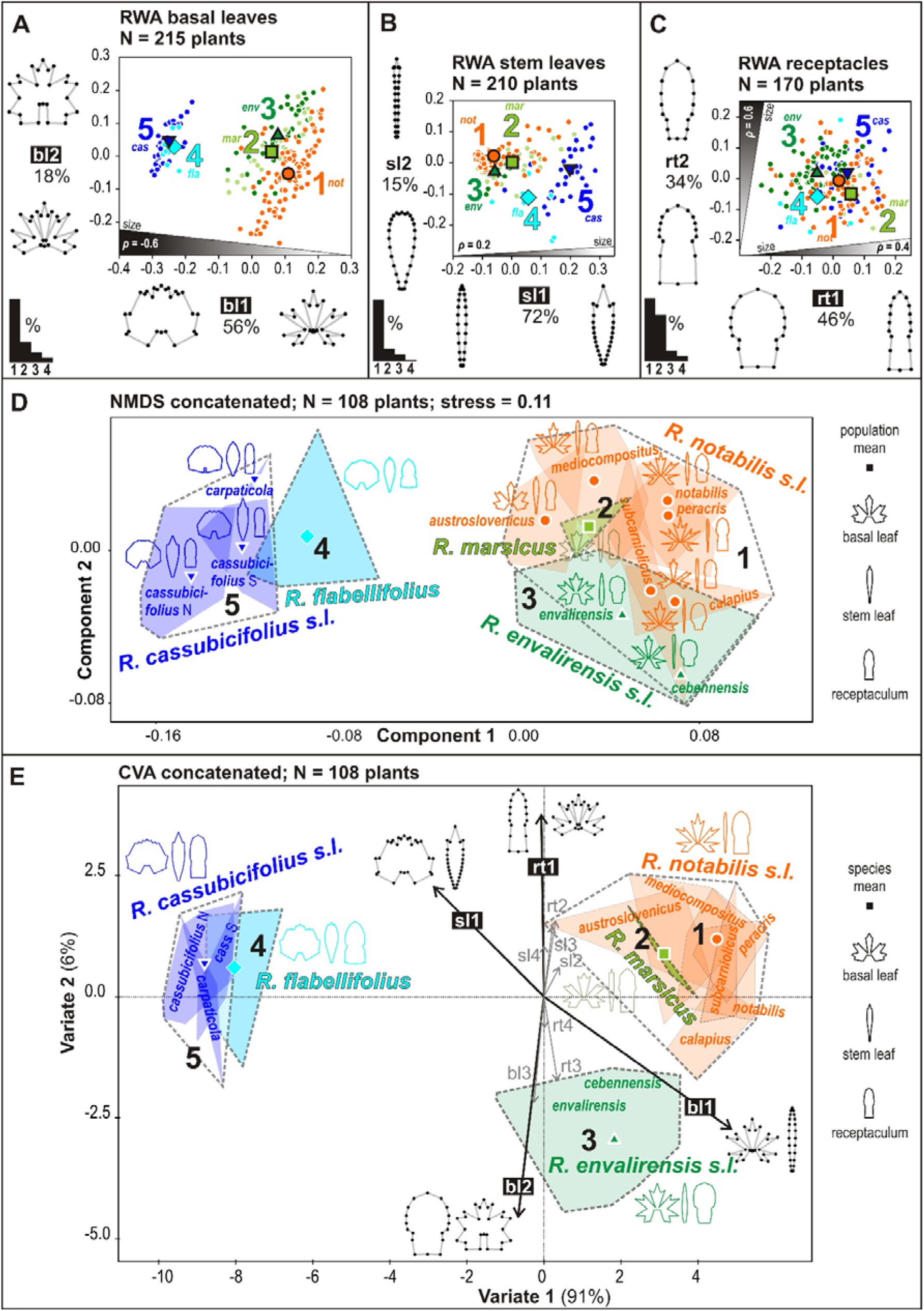
Morphological variation among species. A–C Geometric morphometric analysis of basal leaves (A), stem leaves (B) and receptacles (C) based on relative warps analysis (RWA). Ordination plots of the first and second relative warps show different species clustering according to the three morphological characters. Percentages give variation explained by relative warps, and bar charts in the lower-left corners show the decrease in the explained variation by four major relative warps. The presence of significant positive or negative allometry is given as Spearman’s rho (ρ) value in shaded areas close to the associated relative warps. The area width reflects the correlation strength. Landmark configurations close to relative warps illustrate shape changes. The five accepted species are coded as follows: *R. notabilis* s.l. = 1, red circle; *R. marsicus* = 2, bright-green square; *R. envalirensis* s.l. = 3, dark-green lower triangle; *R. flabellifolius* = 4, bright-blue diamond; *R. cassubicifolius* s.l. – 5, dark-blue upper triangle. d) (D) Morphological differentiation among species based on non-metric multidimensional scaling (NMDS) of the concatenated dataset. Convex hulls delimitate intraspecific morphological variation and symbols show the position of population centroids. Silhouettes close to population centroids illustrate population mean shapes of all three morphological characters. (E) Canonical variates analysis (CVA) highlight shape variables best differentiating among species. Shape variables correspond to relative warp scores and the most important vectors (bl1, bl2, rt1, sl1) correspond to relative warps shown above (A–C).

### Geometric morphometrics - Morphological clustering

Concatenated morphometric data sets containing shape variables extracted from basal leaves, stem leaves and receptacles enabled better resolution of species differences. Non-metric multidimensional scaling analysis of 11 shape variables supported the deep morphological split between *R. cassubicifolius* s.l. on the one side and *R. envalirensis*, *R. marsicus* and *R. notabilis* s.l. on the other side (axis 1; Fig. 6D). This morphological split regards not only the most prominent character, i.e., the basal leaves, but also the stem leaves and to a lesser extent the receptacles. Concerning *R. flabellifolius*, the concatenated analysis confirmed its overall higher similarity to *R. cassubicifolius* s.l. than to other species. Moreover, *R. flabellifolius* was morphologically closer to southern than to northern populations of *R. cassubicifolius* (Fig. 6D). The shape of stem leaves and receptacles separated *R. flabellifolius* from *R. cassubicifolius*. The second ordination axis (component 2) separated between species with dissected basal leaves (right part of the NMDS ordination plot; Fig. 6D), i.e., *R. envalirensis* and *R. notabilis* s.l.. The NMDS analysis suggested that the separation between *R. envalirensis* and *R. notabilis* s.l. was mostly due to basal leaves and receptacles. Canonical variates analysis with five a priori determined groups (= species) supported the clustering of *R. envalirensis* populations apart from *R. notabilis* s.l. populations with main separating characters being the receptacles (rt1) and basal leaves (bl2) (Fig. 6E). *Ranunculus notabilis* exhibited narrower and bigger receptacles than *R. envalirensis* and basal leaves finer divided and with a wider blade base than *R. envalirensis*. *Ranunculus marsicus* possess a combination of traits: the basal leaves were *R. envalirensis*-like, the stem leaves resembled particular populations of *R. notabilis* s.l. with less dissected leaves (*R. austroslovenicus, R. mediocompositus*) and the receptacles were generally *R. notabilis* s.l.-like. Non-parametric multivariate analysis of variance (NP-MANOVA) revealed significant morphological differentiation among *R. cassubicifolius*, *R. envalirensis* and *R. notabilis* with overall (p < 0.001) and pairwise comparisons (p_cas/env_ < 0.001, p_cas/not_ < 0.001 and p_env/not_ < 0.001). *Ranunculus flabellifolius* and *R. marsicus* were not included here due to low sample sizes.

### Indumentum of receptacle

Trichome density of the indumentum of receptacles varied considerably within and among taxa, but it significantly differed among described taxa and accepted taxa (Chi² = 151.69, df = 9, p < 0.001; Chi² = 129.97, df = 4, p < 0.001; see Supplementary Fig. 5 and Table 6). *R. cassubicifolius* s.l. revealed a significant higher trichome density (12.12 / 0.25 mm²) compared to the other species. Whereas *R. marsicus* is completely glabrous and *R. flabellifolius* showed almost no trichomes (0 and 0.22 / 0.25 mm²), *R. envalirensis* and *R. notabilis* s.l. (1.01 and 2.25 / 0.25 mm²) exhibited low receptacle trichome densities.

*Ranunculus notabilis* s.str. was the only taxon of the Illyrian clade that showed a remarkable trichome density (9.50 / 0.25 mm²), not significantly different to *R. cassubicifolius* s.l. (see Supplementary Fig. 5A and Table 6A).A

## Discussion

Here, we present the first comprehensive phylogenomic and geometric morphometric analysis of all hitherto known sexual species of the Eurasian *R. auricomus* complex. The delimitation of sexual species and reconstruction of a phylogenetic framework of progenitors is an important step for the classification of apomictic polyploid complexes (Grant, 1981; Burgess & al., 2015; Hörandl, 2018). We ascertained previously reported ploidy levels and sexual mode of reproduction for all taxa by using flow cytometric methods. In general, we observed congruent patterns among ML, genetic distance in the neighbor-net network and STRUCTURE analyses based on RADseq data (Figs. 3, 4). As an independent test, the coalescent-based species delimitation implemented in STACEY also supported scenarios with a reduced number of species. We investigated five main lineages/clusters that in some cases do not match described species. Six well-defined clusters are easily recognizable in the similarity matrix (Fig. 5). Of these lineages, five could be also characterized by morphometric data and geographical distribution, and represent species in an evolutionary sense (*R. cassubicifolius* s.l.*, R. envalirensis* s.l., *R. flabellifolius, R. marsicus,* and *R. notabilis* s.l.). For species delimitation, a combined workflow of phylogenomic analyses (Fig. 2), based on independent methods (ML, distance and structure analyses, coalescent-based species delimitation) and geometric morphometrics provide a comprehensive picture of genetic and phenotypic differentiation among *R. auricomus* species.

### Phylogenomic data

Phylogenomic analyses based on RADseq and target enrichment proved to be successful in disentangling phylogenetic relationships among sexual species in the less than one million-year-old *R. auricomus* complex. The herein presented comprehensive workflow is applicable to other evolutionary young species complexes that are shaped by low genetic divergence, hybrid origins, introgression and incomplete lineage sorting (ILS).

Both methods were able to resolve distinct evolutionary lineages in the sexual taxa of the *R. auricomus* complex. Results are in line with literature demonstrating that RADseq is able to unravel relationships among populations, closely related plant species and genera more than 50 Mya years old (Baird & al., 2008; Hipp & al., 2014; Cavender-Bares & al., 2015; Tripp & al., 2017; Wagner & al., 2018; Pätzold & al., 2019). RADseq parameter optimization mainly aimed at the assembly of homologous loci and maximization of phylogenetic information in concatenated ML trees resulting in well-resolved phylogeny of the evolutionary young sexual species within the *R. auricomus* complex (see Supporting Figs. 2). However, *de novo* assembly of RADseq reads remains methodically challenging considering various genomic divergence levels within and among individuals/species (see e.g., Guillardín-Calvo & al., 2019; Pätzold & al., 2019; Rancilhac & al., 2019).

Target enrichment of nuclear genes demonstrated its usefulness in resolving phylogenetic relationships above the species level, producing results comparable to those of RADseq and a well-resolved backbone of the tree topology. Phylogenetic reconstruction below species level loses power as demonstrated by the low bootstrap values obtained by the branches within the Illyrian clade. Target enrichment has in several studies shown its usefulness for resolving phylogenetic relationships above genus and family level (Mandel & al., 2014; Sass & al., 2016; Wanke & al., 2017; Vatanparast & al., 2018), and only in a few cases it has been employed on recently radiating species groups (Stephens & al., 2015) or below the species level (Holliday & al., 2016). In *Ranunculus*, target enrichment of nuclear genes is also clearly superior over plastid markers for resolution of species-level relationships (e.g., Emadzade & al., 2015).

In general, we found congruent patterns among ML, genetic distance in the neighbor-net network and STRUCTURE analyses based on RADseq data (Figs. 3, 4). The fully resolved and well-supported phylogeny (BT ∼ 90 - 100) with fully informative nodes is the first one concerning all yet known sexual species within the *R. auricomus* complex. We observed the deepest split in the ML tree, the largest genetic distance in the neighbor-net network and the most distinct genetic structuring between *R. cassubicifolius* s.l. with non-dissected basal leaves and all other species characterized by non-dissected and dissected basal leaves. Previous studies concerning isoenzyme and ITS marker supported the observed split between the most morphological diverged species *R. cassubicifolius* s.l. and *R. notabilis* s.l. (Hörandl, 2004; Hodač & al., 2014). Furthermore, the presence of the non-dissected basal leaf species *R. flabellifolius* from the Western Romanian Carpathians in the clade of the other species with dissected leaves clearly rejects the old morphological Linnean classification of the complex in *R. auricomus* L. s.l. with only dissected basal leaves and *R. cassubicus* L. s.l. with only non-dissected basal leaves (Linnaeus, 1753; but see Kvist, 1987). All described species are monophyletic in the ML tree but ML tree topology, neighbor-net network and genetic structure indicate for less genetic lineages/groups than described species (Figs. 3, 4).

The clade of *R. cassubicifolius* s.l. in the ML tree is obviously genetically distant to the other species in the neighbor-net network possessing an own partition with a geographical E-W gradient in the STRUCTURE analysis (see Fig. 3). BT values and QS scores are high concerning the branch between Southern Carpathian populations and the other species but rather conflicted between the Northern Carpathian and the perialpine populations, suggesting introgression and/or ILS. The high conflict between northern and southern *R. cassubicifolius* populations also introgression (see also Pease et al. (2018) for interpretation of QS scores). The network analysis also supported the presence of reticulation events between northern and southern *R. cassubicifolius* populations. STRUCTURE analyses revealed a northern and a southern cluster of *R. cassubicifolius* s.l. showing small fractions of admixture. Close genetic relationships between these two species were already investigated in Hörandl (2004), Hörandl et al. (2009) and Hodač et al. (2014), fitting to the minor morphological differences in basal and stem leaf traits observed in the present geometric morphometric analyses. Intriguingly, tetraploid *R. cassubicifolius* formed together with diploid samples a specific clade in coalescent-based species tree (Fig. 5; see for details Tomasello et al., subm.). Genetic distance and structure, and the presence of introgression and/or ILS despite geographical barriers within this disjunctly distributed species indicate the presence of one huge, though geographically partitioned, genetic lineage.

*Ranunculus flabellifolius*, *R. envalirensis* s.l. and *R. marsicus*, the closest species of the second main clade in the ML tree that are locally distributed in European mountains (Figs. 1, 3), are genetically far distant from *R. cassubicifolius* s.l. and genetically close to the other Illyrian species (Fig. 4). Whereas the split between *R. flabellifolius* and the other species is fully-supported (BT = 100), the clade formed by *R. envalirensis* s.l. and *R. marsicus* received decreased support (BT = 94 and 69) in both the RADseq based ML tree (see also the presence of discordant patterns in Fig. 3) and in the coalescent-based species tree (Fig. 5). Neighbor-net network and partitioning of genetic clusters in STRUCTURE analysis indicate three genetic clusters (*R. envalirensis* s.str. together with *R. cebennensis*) composed of different subclusters (except *R. envalirensis*) suggesting past reticulation events, i.e., hybrid origins and ILS, or ancient polymorphisms. QD node scores of 0.5 and 0.45 indicate both a more frequent and less frequent alternative topology suggesting ILS, but also the possibility of hybridization with subsequent sequence evolution cannot be excluded. Despite full BT support with an absence of discordant patterns characterizing *R. flabellifolius* as a distinct lineage, the admixture in STRUCTURE and the occurrence of broad three-lobed basal spring leaves in *R. flabellifolius* may hint at an ancient reticulate origin. Furthermore, Reichenbach (1832) and Dunkel et al. (2018) observed that *R. flabellifolius* potentially formed hybrids with other species of the *R. auricomus* complex at the ‘locus classicus’. Hence, recent introgression may have influenced the genetic composition of *R. flabellifolius* species although individuals sampled in the present study possessed characteristic leaf cycle and fan-shaped stem leaves. Introgression could also explain the strange pattern observed in the species delimitation analysis, in which a sample of *R. flabellifolius* (Du-25795/02) forms a distinct cluster, well separated from the other two accessions of the species. Even though from ‘locus classicus’ and morphologically identical to the others, this sample might be an introgressed individual. The dwarf Southeastern European species *R. envalirensis* clusters together with the recently described species *R. cebennensis* occurring in the Massif Central. Both species are morphologically similar (except remarkable plant height differences) as already mentioned by Dunkel & al. (2018) and confirmed by our geometric morphometric analysis (Fig. 6). Together they form a clade in the ML tree and a cluster/partition in our analyses. Due to isolation-by-distance (∼ 300 km) that has been frequently observed in many plant species (Heywood, 1991; Ali & al., 2012), the *R. envalirensis* lineage has been probably diverged into geographical isolated, genetically weakly differentiated sublineages as indicated in the neighbor-net network (see Fig. 1).

*Ranunculus marsicus*, the only known sexual tetraploid species without a diploid cytotype, forms a particular clade in the ML tree and a distinct cluster in the neighbor-net network. Topological conflict and low support of branches involving this species suggest an allopolyploid origin. Genetic structure analysis indicates that *R. envalirensis* s.l., *R. flabellifolius*, a yet unknown species (Fig. 3; cyan partitions) or even an Illyrian species may have contributed to the origin of *R. marsicus*. Interestingly, we did not observe this genetic partition of the yet unknown species in *R. cassubicifolius* s.l. (K3 and K4 of the total data set find also this partition; Supplementary Fig. 7). Results from the coalescent species delimitation analysis indicate a possible contribution of the Illyrian clade to a putative allopolyploid origin of *R. marsicus*, as demonstrated by the moderately high posterior probability of belonging to the same genetic cluster shown by samples from both clades (Fig. 5). *Ranunculus marsicus* and *R. envalirensis* s.l. are the only lineages of the *R. auricomus* complex adapted to subalpine environments. Contact zones between members of the Illyrian group and the other progenitors of a putative allopolyploid *R. marsicus* might have occurred in the Pleistocene due to distributional range shift towards south (of an Illyrian progenitor) and/or expansion (of the putative subalpine progenitor) during cold periods (see Feliner, 2011; Hewitt, 2011). A potential current or past present diploid species in Central Apennines (Dunkel & al., 2018) may have also been a progenitor of *R. marsicus*.

Masci et al. (1994) observed a diploid behavior, i.e., predominantly two alleles per isoenzyme locus in sexual *R. marsicus*, potentially suggesting allopolyploid origin for *R. marsicus*. However, during the filtering processes, only two alleles were allowed removing loci with more than two alleles and potentially biasing allelic diversity of *R. marsicus*. Further research is needed and the application of species network reconstruction approaches (e.g., Yu & Nakhleh, 2015; Oberprieler & al., 2017; Solís-Lemus & al., 2017; Wen & Nakhleh, 2018) and a biogeographical analysis (Tomasello et al., in prep.) might help to unequivocally shed light on the evolution of this tetraploid species.

Regarding the Illyrian group that comprises *R. notabilis* and five recently described species (Dunkel & al., 2018), bootstrap support is decreased whereas discordant patterns remarkably increased in the ML tree (Fig. 3). Genetic distance is low among these species sharing one characteristic genetic partition in varying proportions (orange part; Fig. 3, 4). Discordant tree topologies and the lack of species-specific partitions suggest the presence of introgression and/or ILS. Species of the Illyrian species group, except *R. notabilis*, occur in close neighborhood (see Figure 1; sometimes four species nearby (1 km²), see Dunkel et al. (2018). The absence of crossing barriers apparently allows ongoing gene flow. The genetic differentiation process of the Illyrian clade, which probably started 200,000 years ago and lasts until today, might have caused the observed ILS pattern (see Tomasello et al., subm.). Overall, the Illyrian species group of the *R. auricomus* complex represents one genetic lineage that is weakly geographically structured.

### Geometric morphometrics

Our results quantified the total phenotypic variation concerning basal and stem leaves, and receptacles exhibited by the five sexual species of the *R. auricomus* complex. In accordance with the phylogenomic data, morphometrics pointed out clear morphological clusters corresponding to *R. cassubicifolius* s.l., *R. envalirensis* s.l., and *R. notabilis* s.l. Within the latter, results revealed a morphological differentiation between the most distant morphotypes *R. austroslovenicus* (broad three-dissected leaves) and *R. peracris* (five-dissected leaves) but all other described species as intermediates, indicating population structure of a highly variable species rather than morphologically and genetically well-separated taxa.

The three morphological clusters could be distinguished after combining information from basal leaves, stem leaves and receptacles. When analyzed separately, the morphological characters exhibited only limited species resolution due to overall high phenotypic variation (data not shown). The other two species, *R. flabellifolius* and *R. marsicus,* occupy small restricted distribution areas with only a few known populations, so that their morphological variation is low, particularly for *R. marsicus*. *R. flabellifolius* is morphologically close to *R. cassubicifolius* s.l. due to non-dissected basal leaves, whereas *R. marsicus* is more similar to *R. notabilis* s.l. due to roundish three-lobed basal leaves and linear, sometimes dissected stem leaves in the concatenated analysis.

In general, leaf shapes provided better overall taxonomical resolution than receptacles. In concordance with morphological studies (e.g., Borchers-Kolb, 1983, 1985; Hörandl, 2002; Hörandl & al., 2009), geometric morphometrics confirmed that shape of basal spring leaves discriminates best among the sexual species in the *R. auricomus* complex. Non-dissected basal leaves were expectedly bigger in size than finely dissected phenotypes (particularly within *R. notabilis* s.l.). Apart from the most prominent shape gradient distinguishing *R. cassubicifolius* s.l. from *R. envalirensis* s.l./*R. notabilis* s.l., the basal leaves showed further levels of finer and size-independent shape variation. These minor shape vectors possibly reflect intraspecific variation among populations belonging to the species *R. notabilis* s.l. The total variation of basal leaves extracted from all five sexual species together did not exceed the variation revealed in experimental crossings between *R. cassubicifolius* s.l. and *R. notabilis* s.l. (Hodač & al., 2018). Moreover, the first three major morphological gradients observed in the five sexual species (83% of the total variability) were identical with those revealed in the crossing study by Hodač & al. (2018). However, the genetic structure would not support a hypothesis of a hybrid origin of the species with ‘intermediate’ leaf morphotypes (*R. envalirensis, R. marsicus* and *R. flabellifolius*), from the *R. cassubicifolius* and *R. notabilis* lineages. Instead, it seems that the whole spectrum of leaf shapes is already present in progenitor species.

The major morphological trend in stem leaves distinguished between *R. envalirensis* s.l./*R. notabilis* s.1. and *R. cassubicifolius* s.l. The gradient from small narrow non-dissected stem leaf segments towards broader and dissected ones has also shown a significant covariation with distinct basal leaf shape changes. The outstanding and species-specific fan-shaped stem leaf of *R. flabellifolius* distinguished this species from the other four species. The associated morphological trend was independent of leaf size but again associated with basal leaf variation. Despite less basal and stem leaf differentiation to *R. notabilis* s.l. in geometric morphometric analyses, the subalpine *R. envalirensis* and *R. marsicus* have a remarkably lower plant height (5-15 cm) compared the lowland Illyrian species (10-30 cm). We observed that *R. envalirensis* and *R. marsicus* keep their dwarfism that is typical for alpine plants (Körner, 2003) also under garden culture (2017-2019). In *R. envalirensis* this tiny habit was kept even in garden offspring raised from seeds (pers. obs. of authors).

Interestingly, although basal and stem leaves covariate in their shape changes, they show different morphological gradients. Although the stem leaf middle segments are homologous to middle segments of basal leaves, however, we did not find any relationship between the variance of these structures. Furthermore, trichome density of receptacle remarkably varied within and among species, but discriminated *R. cassubicifolius* s.l. with remarkably higher trichome density from the other accepted species. However, receptacles of the Illyrian species were not glabrous as described by Dunkel et al. (2018). Variation of the indumentum of the receptacle is in general much higher than it was described in previous taxonomic studies (Grau, 1984; Hörandl & Gutermann, 1998c; Dunkel & al., 2018).

From the three morphological shape characters under study, receptacles showed the lowest discriminating power among species and its major shape changes were significantly affected by size variation. This allometry resulted in a general gradient from bigger receptacles (with mostly elongated gynoclinium plus narrow androclinium) towards smaller receptacles (with mostly spherical gynoclinium plus wider androclinium). As mentioned above, variation in receptacles had a limited taxonomic resolution when analyzed separately from other traits. However, distinct vectors of receptaculum variation showed significant covariation with shape changes in other morphological characters. For example, a size-independent morphological trend from receptacles with elongate gynoclinium (plus wide androclinium) towards receptacles with spherical gynoclinium (plus narrow androclinium) covaried with shape changes of basal and stem leaves. In the Nordic species of the complex, characters of the receptacle were regarded as fairly constant, but their variation is also influenced by different environmental conditions (Ericsson, 2001). Phenotypic plasticity of all morphological characters, however, needs to be studied.

### Species delimitation

In general, results of the present study raised the need for a re-classification of the described sexual species within the *R. auricomus* complex. On the one hand, an evolutionary ancestor-descendant lineage concept, including evolutionary role and circumscription of lineages, should be preferred for species delimitation from the conceptual point of view (De Queiroz, 2007; Freudenstein & al., 2017; Sukumaran & Knowles, 2017; Hörandl, 2018). On the other hand, the genetic cluster concept, where a species is defined as a ‘morphologically or genetically distinguishable group of individuals that has few or no intermediates when in contact with other such clusters’ (Mallet, 1995), is also well-applicable when considering the present distance-based genetic and morphometric analyses. A clustering criterion helps specifically for the recognition of distinct morphological entities in groups with a high phenotypic variation and lack of exclusive diagnostic characters, as we observe it in the *R. auricomus* complex. Combinations of leaf and fruit characters rather than single diagnostic features characterize also species within the whole genus *Ranunculus* (Hörandl & Emadzade, 2012). The biological species concept (Mayr, 1942), requiring crossing barriers, is partly supported by the strongly reduced fertility of hybrids between *R. cassubicifolius* s.l. and *R. notabilis* s.l., i.e., the genetically most distant lineages (Hörandl, 2008). Disturbances of meiosis, megasporogenesis and pollen formation in these hybrids are possible triggers for apomixis (Hojsgaard & al., 2014; Barke & al., 2018; Barke et al., subm.). Interestingly, experimental crossings of the Illyrian taxa revealed a higher seed set in interspecific than in intraspecific crosses, supporting our conclusion to merge them all into one species *R. notabilis* s.l. (Rahmsdorf, 2019; Karbstein et al., in prep.). Some tendencies to self-fertility in the Slovenian populations might result in local lineages and morphotypes (Rahmsdorf, 2019; Karbstein et al., in prep.). A rare occurrence of asexual seeds in single individuals of otherwise diploid sexual taxa does not justify a separate taxonomic recognition (Hörandl, 2018). The appearance of rare, asexual seed formation within diploid sexual populations has been observed in other species as well (Schinkel & al., 2016) and may be due to environmental influence (Klatt & al., 2018).

Geographical and altitudinal isolation are the most important factors separating the five main species from each other: *R. cassubicifolius* s.l. in the N and SE edges of the Alps and the Carpathians; *R. flabellifolius* restricted to the Western Romanian Carpathians and Serbia; *R. envalirensis* s.l. to the Massif Central and SE Pyrenees; *R. marsicus* to the Central Apennines; and *R. notabilis* s.l. to the Illyrian region (Figure 1). We suppose that allopatric speciation has shaped the evolution of species. Geographical isolation may have further led to genetic differentiation between the Southern Carpathian and the Northern perialpine populations of *R. cassubicifolius* s.l. and between Pyrenees and Massif Central within *R. envalirensis* s.l.. Low amounts of self-compatibility may have resulted in local differentiation of *R. notabilis* s.l. populations. It might be useful to recognize geographical/ecological entities as subspecies, but more data are needed on ecology and population genetics of the respective taxa. Considering polyploid *R. marsicus*, both ploidy level and geographical isolation separate it from the other species.

We, therefore, propose a new circumscription for the sexual species of the *R. auricomus* complex by rejecting the old classification with purely morphological descriptions and by applying an evolutionary lineage concept based on phylogenetic trees combined with a cluster concept based on distinct genetic and morphological groups. Following this approach, we would have three options of lumping versus splitting of species: We could define (1) two main genetic or morphological lineages/clusters, that are not congruent to each other due the variable position of *R. flabellifolius*; (2) five genetic geographically isolated lineages/clusters with congruent results in phylogenomic and geometric morphometric data, neglecting particular weak genetic and morphological (or local, ecotypic) differentiation; or (3) twelve, though monophyletic, lineages lacking genetic and morphological divergence. We found the best congruence of phylogenomic and morphological data with five distinct lineages/clusters representing species (see Supplementary Fig. 3-6).

Therefore, we accept five, morphologically differentiated, geographically (and altitudinal) well-separated, genetic main lineages as species, to be named after the rules ICBN 2017 (Turland & al., 2017): *R. cassubicifolius* s.l. (incl. *R. carpaticola*), *R. flabellifolius*, *R. envalirensis* s.l. (incl. *R. cebennensis*), *R. marsicus* and *R. notabilis* s.l. (incl. *R. austroslovenicus*, *R. calapius*, *R. mediocompositus, R. peracris* and *R. subcarniolicus.* These five taxa will provide also a robust phylogenetic framework for the analysis of origin and evolution of their apomictic derivatives (Hörandl, 2018).

## Taxonomic Treatment

- *R. flabellifolius* Heuff. ex Rchb., in Fl. Germ. Excurs.: 723,1832. Lectotype: Romania, Banat, ‘in dumetis sylvisque montius [sic!] calc. Banatus’, Apr, May c. 1830, leg. J.A. Heuffel JE00007661 (!). -Syn.: *Ranunculus cassubicus* var. *flabellifolius* (Heuffel ex Rchb.) Borza, *R. cassubicus* auct. europ. p.p. (e.g., Jalas & Suominen, 1989; Tutin & Akeroyd, 1993) non L. (see Kvist, 1987). Distr.: Western Romania, Serbia (Dunkel & al., 2018; this paper, Table 1).
- *R. marsicus* Guss. & Ten., in Florae Neapolitanae Syllogem Appendix quarta: 24,1835. *R. marsicus* var. *marsicus* sensu Dunkel (2011). Type: [Italy, Abruzzo, province of ĹAquila], Piano de 5miglia, leg. M. Tenore, NAP (!) (lectotype, see Dunkel, 2011). Distr.: Central Italy (Masci & al., 1994; Dunkel, 2011); this paper, Table 1). Dunkel (2011) discriminated three varieties: var. *marsicus*, var. *incisior* Dunkel and var. *approximans* Dunkel, whereby the typical variety represents the tetraploid, sexual populations that we analyzed here. The two other varieties have more deeply divided basal leaves and were interpreted as introgressants or hybrids with other co-occurring species of the complex (Dunkel, 2011). Introgression may also explain the occurrence of apomixis in the higher ploidy levels (this paper).
- *R. cassubicifolius* W.Koch, in Ber. Schweiz. Bot. Ges. 49: 553,1939. Type: Switzerland, Kanton Solothurn: ‘Wald östlich Erlenmoss südöstlich Recherswil längs eines Baches am Waldrand, ca. 465 m’, leg. 03 May 1938 W. Koch 38/56, ZT00035121 (!). *R. cassubicus* auct. europ. p.p. (e.g., Jalas & Suominen, 1989; Tutin & Akeroyd, 1993) non L. (see Kvist, 1987). - Syn.: *R. carpaticola* Soó (1965 emend. Hörandl & al., 2009). Distr.: Austria, Germany, Hungary, Italy, Romania, Slovakia, Slovenia, Switzerland (Borchers-Kolb, 1985; Hörandl & al., 2009; Dunkel, 2010; this paper, Table 1). This species turned out to be more widely distributed and more variable than anticipated by previous authors. An infraspecific classification recognizing its disjunct distribution appears desirable but requires further investigations, including the type area and also the autotetraploid sexual cytotypes in Lower Austria (Hörandl & Greilhuber, 2002). The polyploid hybridogenetic apomictic derivatives were excluded from the diploid sexual species (Hörandl & al., 2009).
- *R. envalirensis* Grau, in Mitt. Bot. Staatssamml. München 20: 14,1984. Type: Andorra, alpine Matten zwischen Soldeu und dem Puerto de Envalira, c. 1900 m, 23 May 1970, Merxmüller & Gleisner, M0008123 (!). - Syn.: *Ranunculus auricomus* subsp. *envalirensis* (Grau) Molero & al.; incl. *R. cebennensis* (Dunkel & al., 2018). Distr.: Andorra, France, Spain (Grau, 1984; Dunkel & al., 2018; this paper, Table 1). Already Landolt in Jalas and Suominen (1989) supposed that the diploid populations in the Eastern Pyrenees and the Massif Central match *R. envalirensis*.
- *R. notabilis* Hörandl and Gutermann, in Phyton (Horn) 37: 268, 1998. Type: Austria, Burgenland, 8964/4 Stremtal, Moschendorfer Wald, am Rand 1.,5 km ENE Strem, 220 m, Feuchtwiese, 01 May 1994, leg. E. Hörandl & W. Gutermann, Hö5613 = Gu27897, holotype WU, isotype WU, Hb Gutermann, Hörandl, M, W (!). - Syn.: *R. austroslovenicus* Dunkel (2018), *R. calapius* Dunkel (2018), *R. mediocompositus* Dunkel (2018), *R. peracris* Dunkel (2018), *R. subcarniolicus* Dunkel (2018). Distr.: Austria, Croatia, Slovenia (Hörandl & Gutermann 1998c; Dunkel & al., 2018), this paper, Table 1).

## Identification key

**1** Basal spring leaves broad, non-dissected. The middle segment of the lowermost stem leaf (broad) lanceolate with a variable number of teeth at the leaf tip, or complete lowermost stem leaf fan-like.

**2**

**1\*** Basal spring leaves three-lobed to dissected. The middle segment of the lowermost stem leaf linear to narrow lanceolate, without teeth at the leaf tip, but sometimes with deep sinuses.

**3**

**2** The middle segment of the lowermost stem leaf only at the basis connected to the other segments, therefore lowermost stem leaf segmented, not fan-like.

Receptacles densely hairy.

***R. cassubicifolius* s.l.**

**2\*** The middle segment of the stem leaf connate with the other segment, therefore lowermost stem leaf fan-like. Receptacles almost glabrous.

***R. flabellifolius***

**3** Basal spring leaves divided to dissected into 3-5 segments. Plants 15-30 cm high. Forest zone of the Illyrian region.

***R. notabilis* s.l.**

**3\*** Basal spring leaves lobed to divided into 3-5 segments. Tiny plants (5)-10-(15) cm high. Subalpine zone of the Apennines, Massif Central and Pyrenees.

**4**

**4** The middle segment of the lowermost stem leaf linear with (sometimes deep) sinuses. Central Apennines

***R. marsicus***

**4\*** The middle segment of the lowermost stem leaf linear without sinuses. Massif Central, Eastern Pyrenees.

***R. envalirensis* s.l.**

## Acknowledgments

We thank the German Research Foundation for project funding (DFG, Ho4395/10-1) to E.H., within the priority program ‘Taxon-Omics: New Approaches for Discovering and Naming Biodiversity’ (SPP 1991). We also acknowledge Jennifer Krüger, Birthe H. Barke and Julius Schmidt for technical help, Claudia Pätzold for bioinformatic support, and Silvia Friedrich for curating the garden plants. We thank the Botanical Museum, University of Oslo (O) for silica-gel material of *R. pygmaeus*.

## Data Accessibility Statement

### Code availability

Exemplary bash and R scripts used in analyses are deposited on the Dryad data repository ().

### Data availability

The authors declare that basic data supporting the findings are available within the Supporting Information. Demultiplexed RADseq reads are stored on National Center for Biotechnology Information Sequence Read Archive (SRA): BioProject ID XXX. Flow cytometric, genetic and morphometric data are available on Dryad data repository

