## Supplementary Fig. 1 for "Phylogenomics supported by geometric morphometrics reveals delimitation of sexual species within the polyploid apomictic *Ranunculus auricomus* complex (Ranunculaceae)"

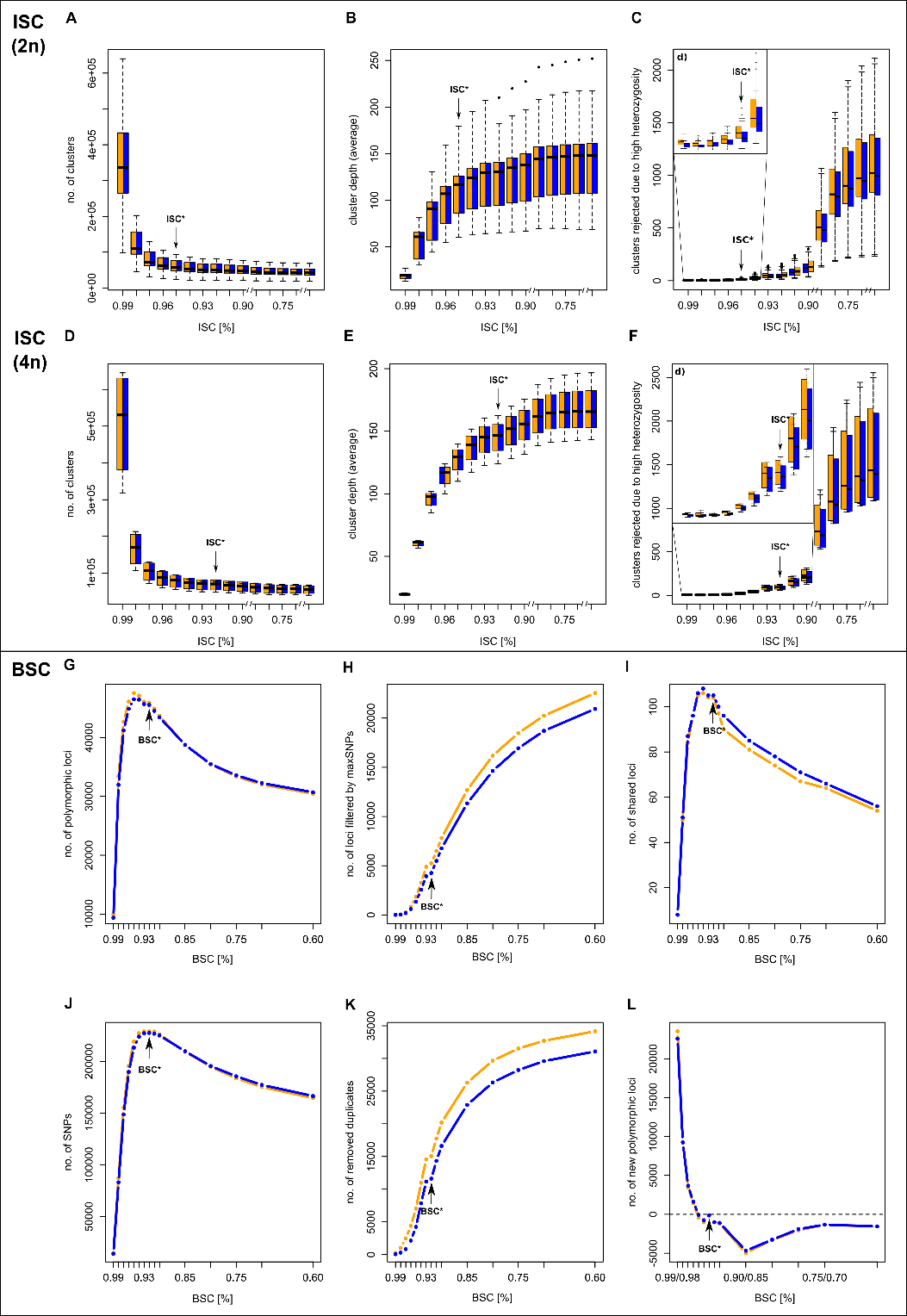

**Supplementary Fig. 1.** Evaluation of in-sample clustering threshold (ISCT) concerning diploid and tetraploid sexual taxa within the *R. auricomus* complex, and between-sample clustering threshold (BSCT) concerning the total dataset. We illustrated boxplots of (A, D) number of clusters, (B, E) cluster depth (average) and (C, F) clusters rejected due to high heterozygosity in relation to ISCT. Due to huge differences within clusters rejected due to high heterozygosity across ICSTs, we added an extended plot of ICSTs ranging from 94 to 99%. We illustrated boxplots of (G) number of polymorphic loci, (H) no. of SNPs and (I) no. of loci filtered by maxSNPs, (J) no. of removed duplicates, (K) no. of shared loci across samples and (L) no. of new polymorphic loci in relation to BSCT. Orange boxplots represent ‘mindepth6’ setting, and blue boxplots refer to ‘mindepth12’ setting. ISCT* = selected ISCT for cluster optimization.
