## Supplementary Fig. 2 for "Phylogenomics supported by geometric morphometrics reveals delimitation of sexual species within the polyploid apomictic *Ranunculus auricomus* complex (Ranunculaceae)"

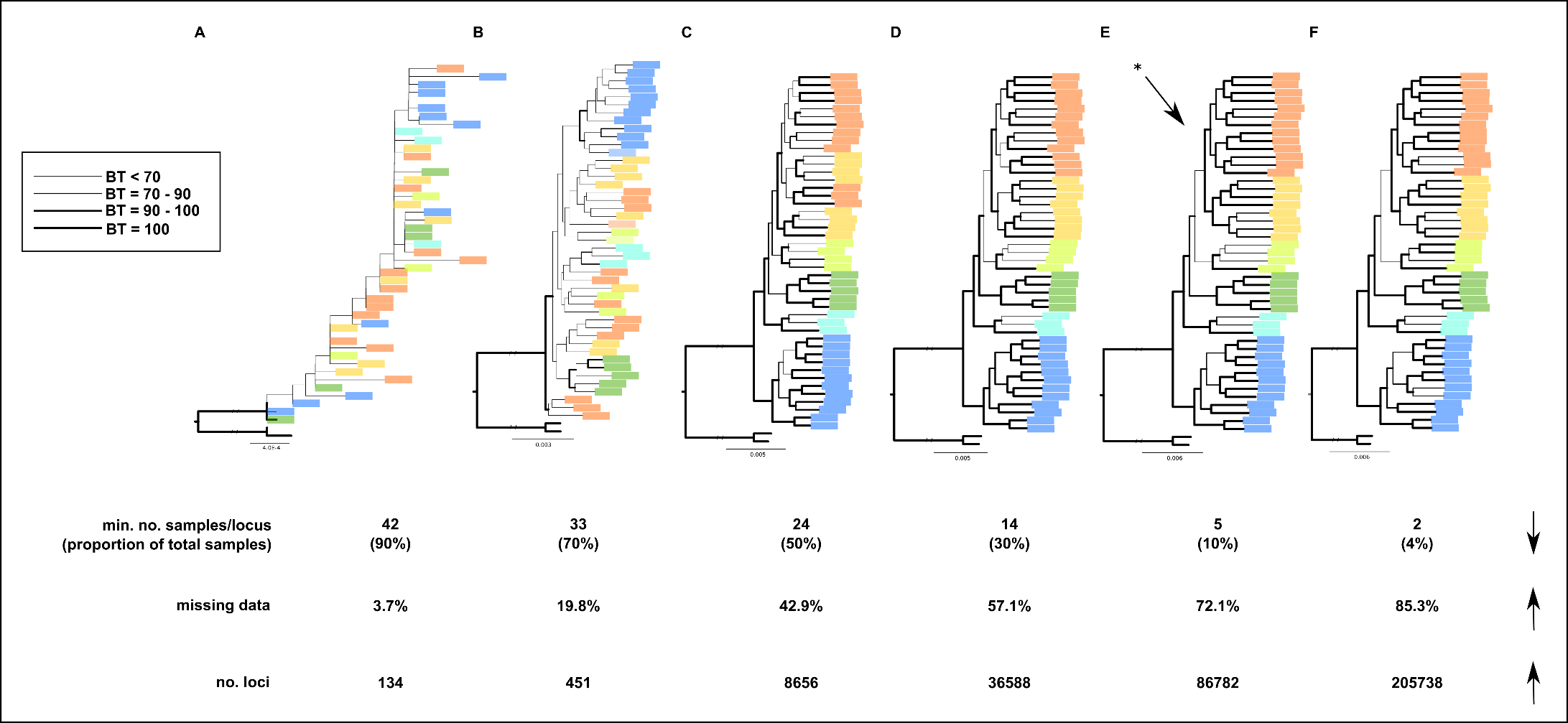

**Supplementary Fig. 2.** Maximum likelihood (ML) trees of different minimum no. samples per locus (2 (4%), 5 (10%), 14 (30%), 24 (50%), 33 (70%) and 42 (90%), A–F). Line strengths represent the magnitude of the bootstrap value concerning the previous node (see legend within the figure). Taxa are colored according to accepted species. Below the ML trees, we put details on minimum no. of samples per locus (and proportion of total samples), percent of missing data, and no. of loci. Arrows specify the trend (either positive or negative).
