## Supplementary Fig. 3 for "Phylogenomics supported by geometric morphometrics reveals delimitation of sexual species within the polyploid apomictic *Ranunculus auricomus* complex (Ranunculaceae)"

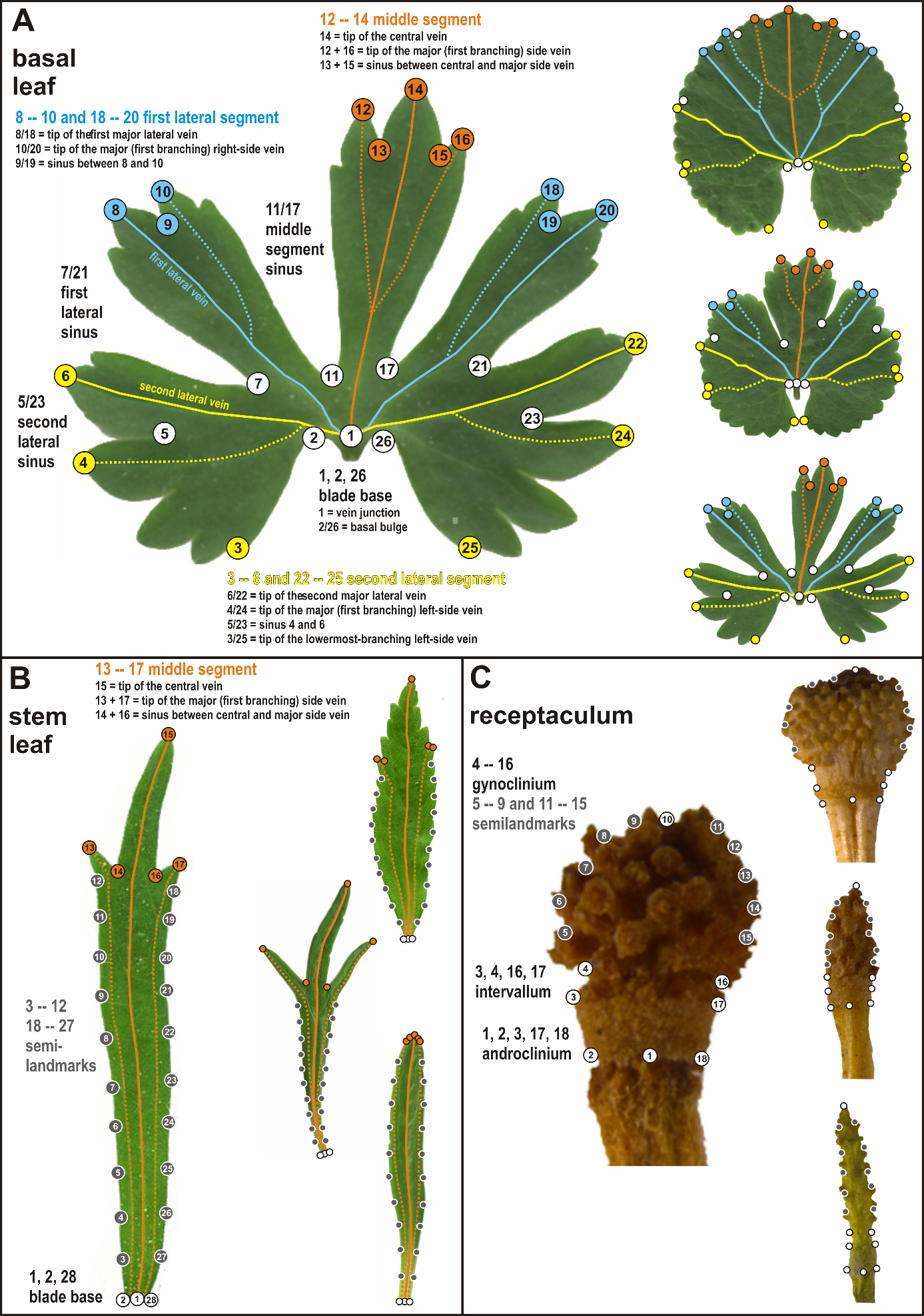

**Supplementary Fig. 3.** Landmark digitization. (A) Delimitation of 26 landmarks on basal leaf outline following homologous leaf venation pattern, segments and sinuses. (B) Delimitation of 8 landmarks (white and orange dots) and 20 semilandmarks (grey dots) on the stem leaf outline. (C) Delimitation of 8 landmarks (white dots) and 10 semilandmarks (grey dots) on the receptacle outline. Images on the right side of each figure illustrate a high morphological variation of species within the *Ranunculus auricomus* complex. The symmetrization of the object left and right halves happen in three steps: (1) exchange of left vs. right object halves, (2) Procrustes superimposition (Zelditch & al., 2012) of original and mirrored objects, (3) averaging of original and corresponding mirrored objects. In the first step, we put original landmark configurations together with their mirrored counterparts into one dataset. The mirrored landmark configurations have been acquired by exchanging x- and y-coordinates within each pair of symmetrically corresponding landmarks. In basal leaves, for example, we reciprocally exchanged the positions of landmarks 2 and 26 with respect to the axis of bilateral symmetry as defined by landmarks 1 and 14. In the final mirrored configurations (after exchanging all corresponding landmarks), we have multiplied all x-coordinates by -1. In the next step, both original and mirrored configurations have been subjected to the Procrustes superimposition (Zelditch & al., 2012). Procrustes fit method translates centroids of all configurations to the same point [0,0] and rotates the configurations so that distances among homologous landmarks are minimized and rescales the configurations to the unit size. Images consisting of landmark configurations without semilandmarks (i.e., basal leaves) have been superimposed in the program MorphoJ (Klingenberg, 2011), and those with semilandmarks (i.e., stem leaves and receptacles) in TpsRelw vers. 1.70 (Rohlf, 2015) allowing semilandmarks (without fixed position) to slide along curves between adjacent (semi-)landmarks. Superimposed landmark configurations, i.e., original configurations and their corresponding mirrored counterparts, have been subsequently averaged resulting in configurations perfectly symmetric with respect to the axis of object bilateral symmetry. Symmetrized landmark configurations representing original leaf and receptacle shapes have been averaged over each plant using the software MorphoJ (Klingenberg, 2011).
