## Supplementary Fig. 4 for "Phylogenomics supported by geometric morphometrics reveals delimitation of sexual species within the polyploid apomictic *Ranunculus auricomus* complex (Ranunculaceae)"

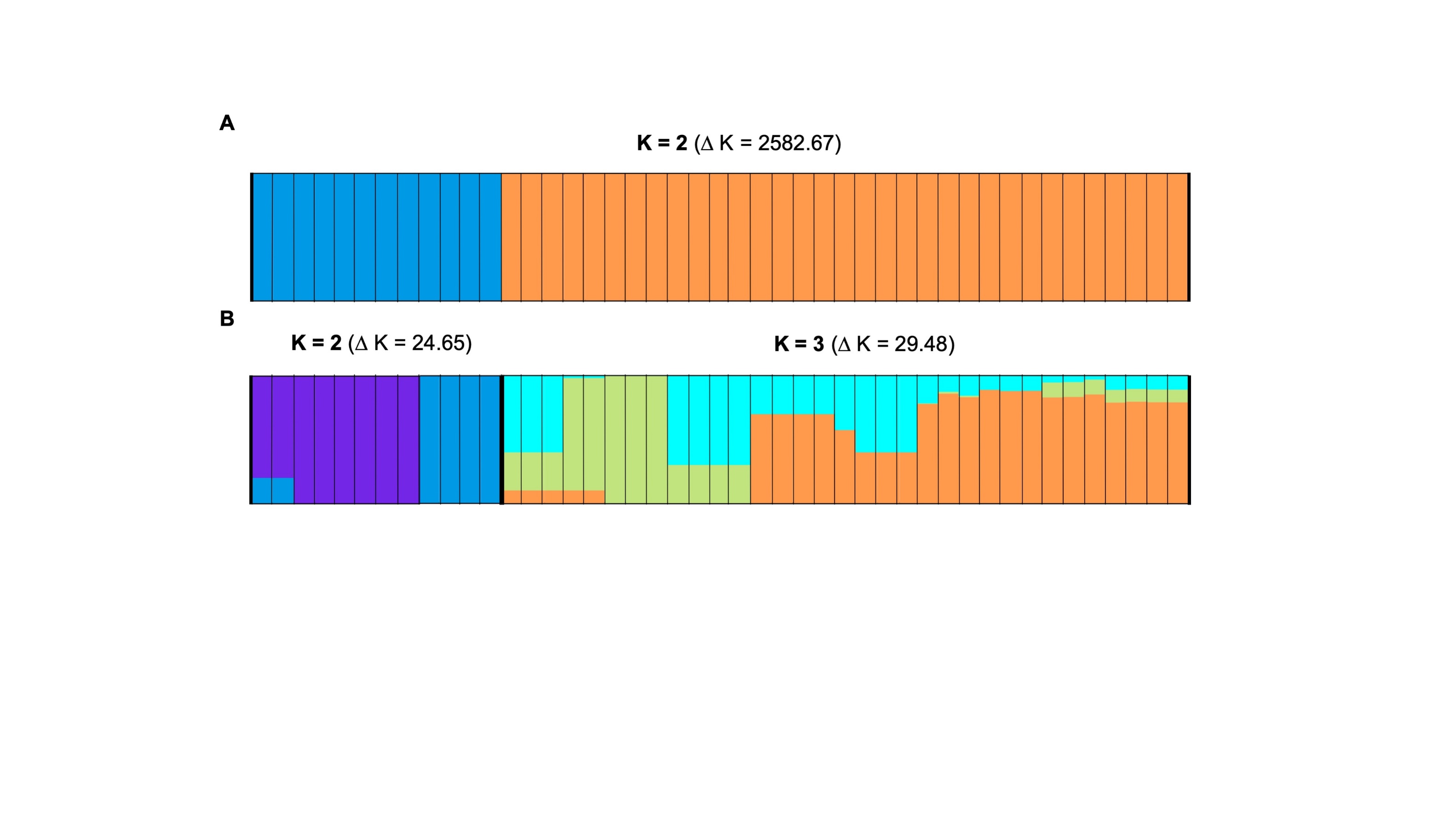

**Supplementary Fig. 4.** Bar graphs of the genetic structure analysis are drawn for (A) the total dataset and (B) for each of the two subsets (see Supplementary Table 5 for results of the Evanno tests). We chose the data set with the likeliest K value.
