## Supplementary Fig. 5 for "Phylogenomics supported by geometric morphometrics reveals delimitation of sexual species within the polyploid apomictic *Ranunculus auricomus* complex (Ranunculaceae)"

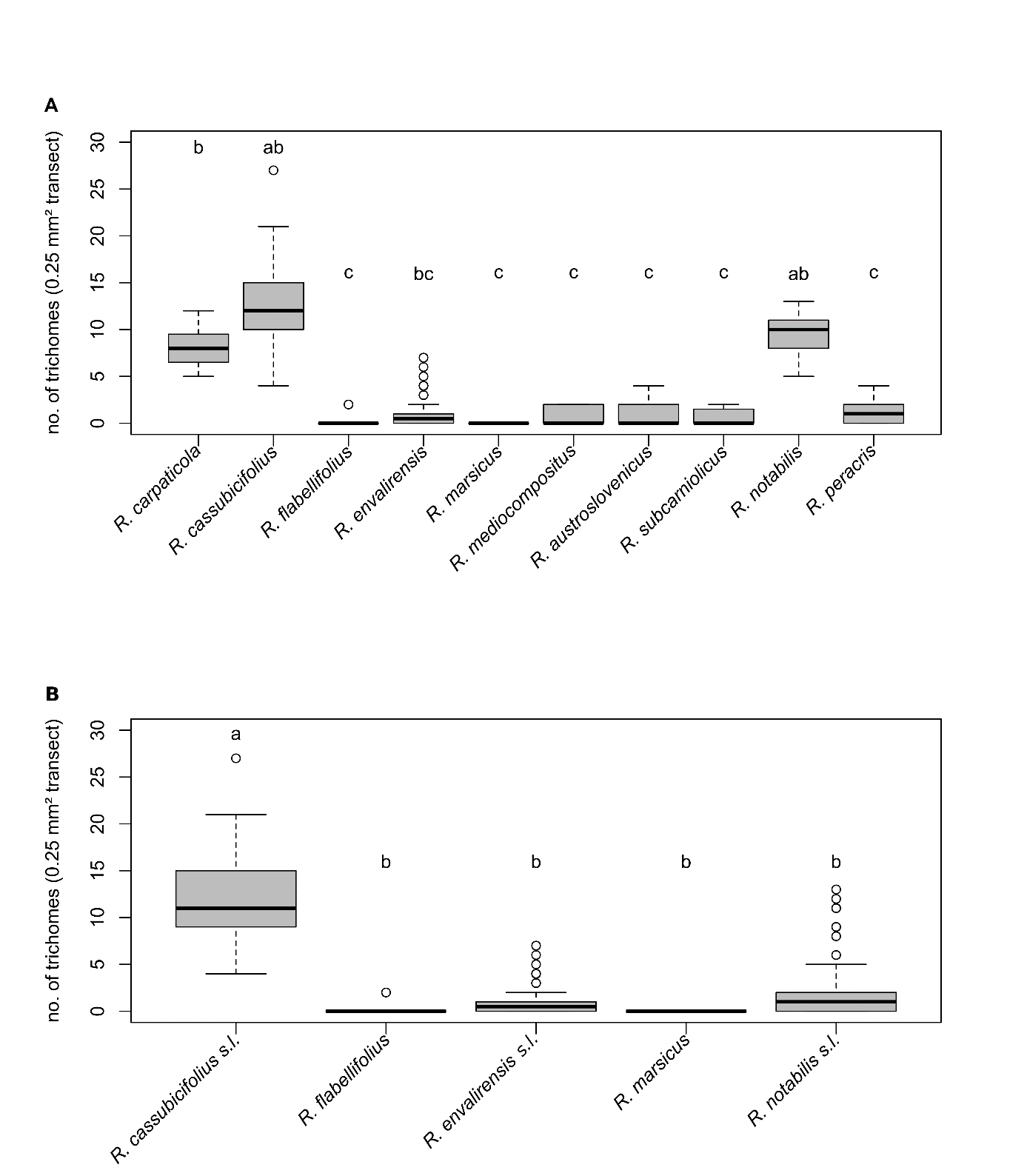

**Supplementary Fig. 5.** Boxplots of receptacle trichome density for (A) all described taxa and (B) all accepted taxa. The number of trichomes on a 0.25 mm² transect. In total, we assessed 219 receptacles. Significant differences in trichome density (A) among species (among all ꭓ² = 151.69, df = 9, p < 0.00) and (B) accepted species among all (ꭓ² = 129.97, df = 4, p < 0.001). Pairwise differences are indicated by superscript letters (see Supplementary Table 6).
