## Supplementary Fig. 6 for "Phylogenomics supported by geometric morphometrics reveals delimitation of sexual species within the polyploid apomictic *Ranunculus auricomus* complex (Ranunculaceae)"

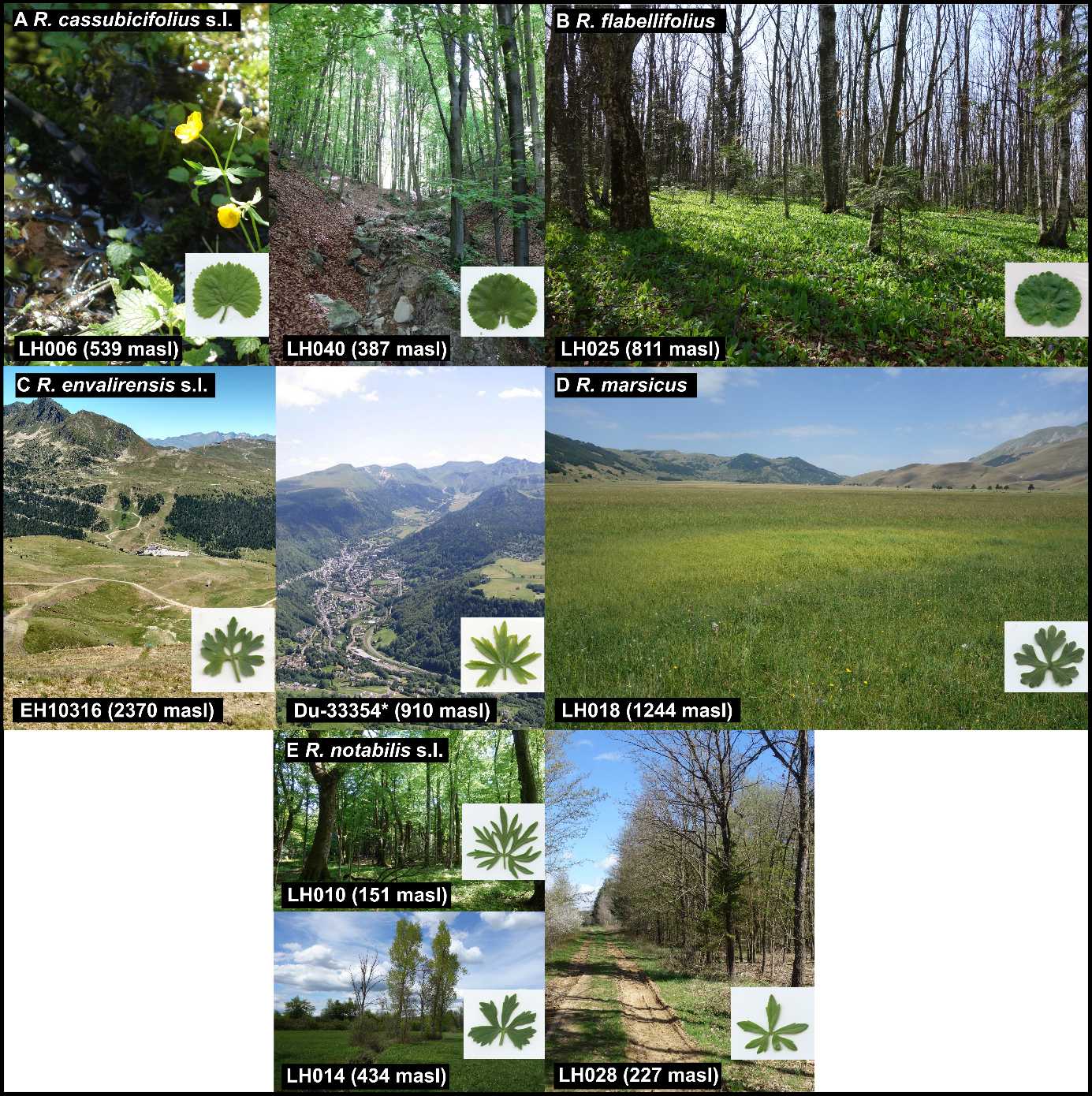

**Supplementary Fig. 6.** Habitats, and basal spring leaf and flower shapes of accepted sexual species within the *R. auricomus* complex (former names in brackets): (A) *Ranunculus cassubicifolius* s.l. inhabits humid beech, hornbeam forests and streamsides (LH006 = *R. cassubicifolius*, LH040 = *R. carpaticola*), (B) *R. flabellifolius* (LH025 = *R. flabellifolius*) was found in forests in a restricted area of the South Carpathians, (C) *R. envalirensis* s.l. (EH10316 = *R. envalirensis*, Du-33354 *R. cebennensis** inhabits (alpine) meadows in Massif Central and Pyrennees, (D) *R. marsicus* (LH018 = *R. marsicus*) occupies humid alpine meadows (three-lobed leaf) and (E) *R. notabilis* s.l. (LH010 = *R. peracris*, LH014 = *R. mediocompositus* and LH028 = *R. notabilis*, *R. austroslovenicus*, *R. subcarniolicus*, *R. calapius* (not illustrated) occurs in deciduous forest, forest and waysides, and humid to marshy meadows. Usually, five corolla leaves characterize the sexual species. We illustrated the most dissected-leaf morphotypes (taxonomically most informative). * The picture on the right shows Mont-Dore near sampling location Du-33354 *R. cebennensis* (edited; source: CC BY-SA 2.0 fr, https://commons.wikimedia.org/w/index.php?curid=211135). masl = meters above sea level.
