## Supplementary Fig. 7 for "Phylogenomics supported by geometric morphometrics reveals delimitation of sexual species within the polyploid apomictic *Ranunculus auricomus* complex (Ranunculaceae)"

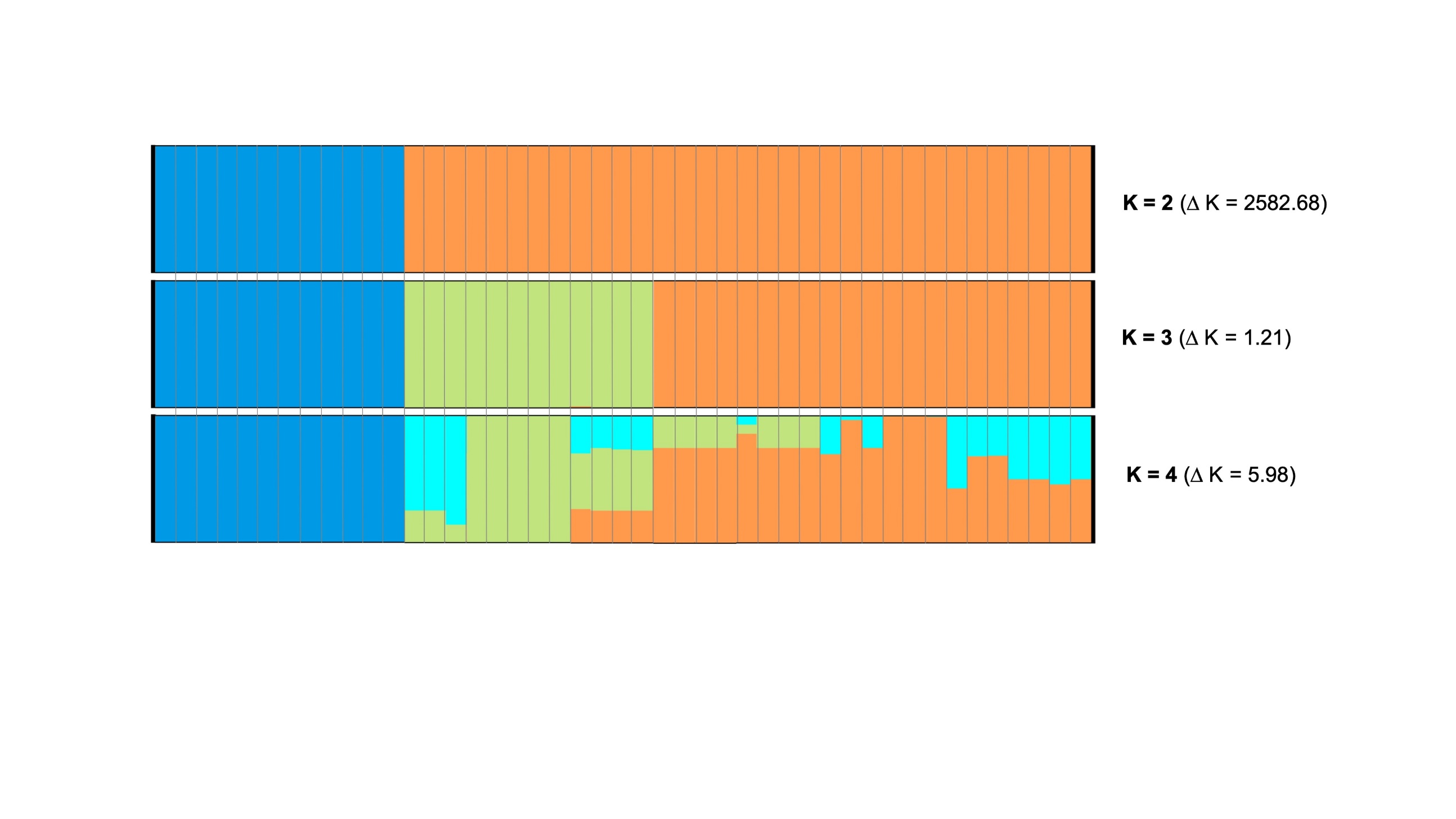

**Supplementary Fig. 7.** Bar graphs of the genetic structure analysis are drawn for the total dataset (see Supplementary Table 5 for results of the Evanno tests). We chose the data set with the likeliest K value, and K = 3 and K = 4
