## Supplementary Table 1 for "Phylogenomics supported by geometric morphometrics reveals delimitation of sexual species within the polyploid apomictic *Ranunculus auricomus* complex (Ranunculaceae)"

**Supplementary Table 1.** Description of the basic flow cytometric procedure. Steps mainly followed Klatt et al. (2016) and Barke et al. (2018).

| **Step** | **Description** |
| --- | --- |
| 0 | We used Otto buffers to first isolate and then stain nuclei (Otto, 1990; Doležel & Bartoš, 2005; Doležel & al., 2007). The composition of buffers followed (Otto, 1990) with slight modifications: 100 ml Otto I buffer are composed of 99 ml ddH_2_O, 2.1014 g citric acid and 0.5 ml Tween 20, and 100 ml Otto II contained 99 ml ddH_2_O, 5.6784 g dibasic sodium phosphate and 1 ml (300µg/ml) 4′,6-Diamidin-2-phenylindol (DAPI) solution. |
| 1 | We put 200 µl extraction buffer Otto I to the ground leaf material and immediately inverted samples for 30 s to isolate the nuclei from the plant cells. |
| 2 | Afterward, we filtered samples through CellTrics® filters with a mesh of 30 µm into flow cytometric sample tubes. |
| 3 | We stained samples with 800 µl Otto II containing DAPI. |
