## Supplementary Table 2 for "Phylogenomics supported by geometric morphometrics reveals delimitation of sexual species within the polyploid apomictic *Ranunculus auricomus* complex (Ranunculaceae)"

**Supplementary Table 2.** Quartet sampling results (Pease & al., 2018) concerning the quartet fidelity (QF) score.

| **taxon** | **QF** |
| --- | --- |
| *R. austroslovenicus* Du-30441/01 SI | 0.82 |
| *R. austroslovenicus* Du-30442/01 SI | 0.87 |
| *R. austroslovenicus* LH012/01 SI | 0.81 |
| *R. austroslovenicus* LH012/08 SI | 0.84 |
| *R. calapius* Du/34889/01 HR | 0.77 |
| *R. calapius* Du/34889/11 HR | 0.69 |
| *R. calapius* Du/35351/15 HR | 0.76 |
| *R. carpaticola* 9126/02 RO | 0.86 |
| *R. carpaticola* 9126/06 RO | 0.86 |
| *R. carpaticola* LH040/04 SK | 0.80 |
| *R. cassubicifolius* Du-15980/01 IT | 0.87 |
| *R. carpaticola* Du-21047/01 HU | 0.79 |
| *R. cassubicifolius* Du-28673/01 SI | 0.85 |
| *R. cassubicifolius* LH006/05 DE | 0.90 |
| *R. cassubicifolius* LH006/17 DE | 0.90 |
| *R. cassubicifolius* LH007/06 AT | 0.92 |
| *R. cassubicifolius* LH007/11 AT | 0.88 |
| *R. cassubicifolius* LH016/01 SI | 0.88 |
| *R. cassubicifolius* LH016/14 SI | 0.86 |
| *R. cebennensis* Du-33354/02 FR | 0.89 |
| *R. cebennensis* Du-33354/21 FR | 0.86 |
| *R. envalirensis* 10316/14 AD | 0.88 |
| *R. envalirensis* Du-29983/01 AD | 0.88 |
| *R. envalirensis* Du-29988/01 FR | 0.91 |
| *R. flabellifolius* Du-25795 RO | 0.80 |
| *R. flabellifolius* LH023/04 RO | 0.82 |
| *R. flabellifolius* LH025/03 RO | 0.80 |
| *R. marsicus* Du-23722/01 IT | 0.82 |
| *R. marsicus* LH017/01 IT | 0.68 |
| *R. marsicus* LH018/02 IT | 0.76 |
| *R. marsicus* LH018/18 IT | 0.71 |
| *R. mediocompositus* LH014/03 SI | 0.72 |
| *R. mediocompositus* LH014 10 SI | 0.76 |
| *R. mediocompositus* LH015/03 SI | 0.70 |
| *R. mediocompositus* LH015 10 SI | 0.72 |
| *R. notabilis* 10137/3 AT | 0.80 |
| *R. notabilis* 10137/8 AT | 0.82 |
| *R. notabilis* LH028/02 AT | 0.80 |
| *R. peracris* LH010/01 SI | 0.85 |
| *R. peracris* LH010/09 SI | 0.87 |
| *R. peracris* LH011/01 SI | 0.81 |
| *R. peracris* LH011/14 SI | 0.82 |
| *R. sceleratus* 10426/1 DE | 0.93 |
| *R. sceleratus* 10426/2 DE | 0.92 |
