## Supplementary Table 3 for "Phylogenomics supported by geometric morphometrics reveals delimitation of sexual species within the polyploid apomictic *Ranunculus auricomus* complex (Ranunculaceae)"

**Supplementary Table 3.** Results of the Evanno test concerning the total dataset, the subset consisting of *R. carpaticola* and *R. cassubicifolius*, and the subset consisting of *R. flabellifolius*, …, and the Illyrian species. We highlighted the likeliest K value. See the documentation of STRUCTURE HARVESTER (Earl & vonHoldt, 2012) for details.

|  | **K value** | **Replicates** | **Mean LnP(K)** | **Stdev LnP(K)** | **Ln'(K)** | **\|Ln''(K)\|** | **Delta K** |
| --- | --- | --- | --- | --- | --- | --- | --- |
| **total dataset** |  |  |  |  |  |  |  |
|  | 1 | 10 | -780507.69 | 487.17 | NA | NA | NA |
|  | 2 | 10 | -667325.97 | 573.46 | 113181.72 | 1481068.94 | 2582.68 |
|  | 3 | 10 | -2035213.19 | 4311638.14 | -1367887.22 | 5198509.93 | 1.21 |
|  | 4 | 10 | -8601610.34 | 16689733.94 | -6566397.15 | 99773314.35 | 5.98 |
|  | 5 | 9 | -114941321.8 | 106168812.4 | -106339711.5 | 75751986.4 | 0.71 |
|  | 6 | 9 | -145529046.9 | 190254761.3 | -30587725.1 | 137716322.8 | 0.72 |
|  | 7 | 9 | -313833094.8 | 306028640.8 | -168304047.9 | 221593315.9 | 0.72 |
|  | 8 | 9 | -260543826.8 | 117831677.5 | 53289268.04 | 108450238.5 | 0.92 |
|  | 9 | 8 | -315704797.2 | 229997413.9 | -55160970.43 | 157638700.7 | 0.69 |
|  | 10 | 8 | -528504468.4 | 484253936.3 | -212799671.2 | 258841562.8 | 0.53 |
|  | 11 | 5 | -482462576.7 | 339927580.2 | 46041891.64 | NA | NA |
| **subet (*carp.*/cass.)** |  |  |  |  |  |  |  |
|  | 1 | 10 | -302819.26 | 213 | NA | NA | NA |
|  | 2 | 10 | -4130631.99 | 1809055.97 | -3827812.73 | 44595959.68 | 24.65 |
|  | 3 | 10 | -52554404.4 | 33638282.3 | -48423772.41 | 19747862.78 | 0.59 |
|  | 4 | 10 | -120726039.6 | 67724624.68 | -68171635.19 | 18384174.67 | 0.27 |
|  | 5 | 10 | -207281849.5 | 128589521.6 | -86555809.86 | 191557765.2 | 1.49 |
|  | 6 | 10 | -485395424.5 | 182136946.1 | -278113575 | 216404468.8 | 1.19 |
|  | 7 | 6 | -547104530.7 | 370028569.7 | -61709106.19 | NA | NA |
| **subset (*flab.*/.../*per.*)** |  |  |  |  |  |  |  |
|  | 1 | 10 | -672385.94 | 274.02 | NA | NA | NA |
|  | 2 | 10 | -1531770.84 | 1810423.94 | -859384.9 | 1038639.82 | 0.57 |
|  | 3 | 10 | -1352515.92 | 1194659.54 | 179254.92 | 35218205.49 | 29.48 |
|  | 4 | 10 | -36391466.49 | 44189080.16 | -35038950.57 | 105634921.9 | 2.39 |
|  | 5 | 10 | -177065338.9 | 78338755.9 | -140673872.4 | 237468525.6 | 3.03 |
|  | 6 | 10 | -555207736.9 | 237071810.9 | -378142398 | 239094479.8 | 1.01 |
|  | 7 | 10 | -694255655.1 | 456645146.8 | -139047918.2 | 406841039.9 | 0.89 |
|  | 8 | 10 | -1240144613 | 507374741.5 | -545888958.2 | 296794947.5 | 0.58 |
|  | 9 | 10 | -1489238624 | 596294321.3 | -249094010.6 | 474518235.4 | 0.8 |
|  | 10 | 5 | -2212850870 | 741454925.4 | -723612246 | NA | NA |
