## Supplementary Table 4 for "Phylogenomics supported by geometric morphometrics reveals delimitation of sexual species within the polyploid apomictic *Ranunculus auricomus* complex (Ranunculaceae)"

**Supplementary Table 4.** Herbarium specimens or data from the literature (in PDF format) incorporated in the geometric morphometric analysis. If a herbarium specimen or data from the literature (PDF) from the locus classicus were included, we marked them with ‘x’. A particular morphological trait is also marked with ‘x’ if it was included in data analysis. See also Table 1 for most location details.

| **Herbarium specimen / PDF** | **Herbarium ID** | **Publication** | **Taxon** | **Locus classicus** | **Basal leave(s)** | **Stem leave(s)** | **Receptacle(s)** |
| --- | --- | --- | --- | --- | --- | --- | --- |
| PDF | Du-25795 | (Dunkel & al., 2018) | *R. flabellifolius* | x | x | x |  |
| herbar specimen | Du-30441 |  | *R. austroslovenicus* | x | x | x | x |
| herbar specimen / PDF | Du-28639 | (Dunkel & al., 2018) | *R. mediocompositus* |  | x | x | x |
| PDF | Du-29983 | (Dunkel & al., 2018) | *R. envalirensis* |  | x | x |  |
| herbar specimen | GOET019892 |  | *R. envalirensis* |  | x | x | x |
| herbar specimen | M-0008124 |  | *R. envalirensis* |  | x | x |  |
| herbar specimen | M-0008125 |  | *R. envalirensis* |  | x | x |  |
| herbar specimen / PDF | Du-30446 | (Dunkel & al., 2018) | *R. peracris* |  | x | x | x |
| herbar specimen / PDF | Du-33354 | (Dunkel & al., 2018) | *R. cebennensis* | x | x | x | x |
| herbar specimen | Du-34772 |  | *R. subcarniolicus* |  | x | x | x |
| PDF | Du-34773 | (Dunkel & al., 2018) | *R. subcarniolicus* |  | x | x |  |
| herbar specimen / PDF | Du-34889 | (Dunkel & al., 2018) | *R. calapius* | x | x | x | x |
| herbar specimen | Du-35342 |  | *R. calapius* |  | **x** | **x** | **x** |
| herbar specimen | Du-35351 |  | *R. calapius* |  | **x** | **x** | **x** |
