## Supplementary Table 5 for "Phylogenomics supported by geometric morphometrics reveals delimitation of sexual species within the polyploid apomictic *Ranunculus auricomus* complex (Ranunculaceae)"

**Supplementary Table 5.** Flow cytometry of leaves (FC) and single seed flow cytometric seed screening (ssFCSS) of sexual diploid and tetraploid species within the *R. auricomus* complex. We only measured populations with yet unknown DNA ploidy and reproductive pathway. References for already published DNA ploidy and reproductive pathway measurements are given in the text. N_ind_ = number of individuals, N_seeds_ = number of seeds.

| **Pop ID** | **Taxon** |  | **N_ind_** |  | **DNA ploidy** |  | **N_ind_** | **N_seeds_** | **Ploidy Embryo** | **Ploidy Endosperm** | **Peak Index (Endosperm : Embryo)** | **Reproductive Pathway** |
| --- | --- | --- | --- | --- | --- | --- | --- | --- | --- | --- | --- | --- |
|  |  | **FC** |  |  |  | **FCSS** |  |  |  |  |  |  |
| *LH012* | *R. austroslovenicus* |  | 11 |  | 2C |  | 3 | 15 | 2C | 3C | 1.53 | sexual |
| *Du-34889* | *R. calapius* |  | 1 |  | 2C |  |  |  |  |  |  |  |
| *Du-35351* | *R. calapius* |  | 1 |  | 2C |  |  |  |  |  |  | sexual |
| *LH040* | *R. carpaticola* |  | 11 |  | 2C |  | 3 | 14 | 2C | 3C | 1.50 | sexual |
| *LH006* | *R. cassubicifolius* |  | 15 |  | 2C |  | 3 | 15 | 2C | 3C | 1.54 | sexual |
| *LH007* | *R. cassubicifolius* |  | 13 |  | 2C |  | 3 | 15 | 2C | 3C | 1.49 | sexual |
| *LH016* | *R. cassubicifolius* |  | 9 |  | 2C |  | 3 | 12  2 | 2C  2C | 3C  6C | 1.54  2.98 | sexual  asexual |
| *Du-33354* | *R. cebennensis* |  | 1 |  | 2C |  |  |  |  |  |  |  |
| *10316* | *R. envalirensis* |  | 36 |  | 2C |  | 3 | 15 | 2C | 3C | 1.50 | sexual |
| *LH023* | *R. flabellifolius* |  | 6 |  | 2C |  | 1 | 5 | 2C | 3C | 1.48 | sexual |
| *LH025* | *R. flabellifolius* |  | 3 |  | 2C |  | 2 | 6 | 2C | 3C | 1.52 | sexual |
| *LH017* | *R. marsicus* |  | 9 |  | 4C, 6C |  | 3 | 1  8 | 4C  6C | 6C  9C | 1.53  2.90 | sexual  asexual |
| *LH018* | *R. marsicus* |  | 14 |  | 4C |  | 3 | 23 | 2C | 3C | 1.47 | sexual |
| *LH014* | *R. mediocompositus* |  | 9 |  | 2C |  | 3 | 15 | 2C | 3C | 1.48 | sexual |
| *LH015* | *R. mediocompositus* |  | 9 |  | 2C |  | 3 | 15 | 2C | 3C | 1.49 | sexual |
| *10137* | *R. notabilis* |  | 2 |  | 2C |  |  |  |  |  |  |  |
| *LH028* | *R. notabilis* |  | 6 |  | 2C |  |  |  |  |  |  |  |
| *LH010* | *R. peracris* |  | 11 |  | 2C |  | 3 | 15 | 2C | 3C | 1.61 | sexual |
| *LH011* | *R. peracris* |  | 15 |  | 2C |  | 3 | 15 | 2C | 3C | 1.50 | sexual |
| *LH013* | *R. subcarniolicus* |  | 9 |  | 2C |  | 3 | 14 | 2C | 3C | 1.52 | sexual |
