## Supplementary Table 6 for "Phylogenomics supported by geometric morphometrics reveals delimitation of sexual species within the polyploid apomictic *Ranunculus auricomus* complex (Ranunculaceae)"

**Supplementary Table 6.** Test statistics for (A) all described taxa and (B) all accepted taxa. Mean trichome density per 0.25 mm² transect of taxa and pairwise p values based on pairwise comparisons using the Wilcoxon rank-sum test with holm correction are given between taxa.

**A**

|  | ***R. carpaticola*** | ***R. cassubicifolius*** | ***R. flabellifolius*** | ***R. envalirensis*** | ***R. marsicus*** | ***R. mediocompositus*** | ***R. austroslovenicus*** | ***R. subcarniolicus*** | ***R. notabilis*** | ***R. peracris*** |
| --- | --- | --- | --- | --- | --- | --- | --- | --- | --- | --- |
| ***R. cassubicifolius*** | 0.18 |  |  |  |  |  |  |  |  |  |
| ***R. flabellifolius*** | 0.01 | <0.001 |  |  |  |  |  |  |  |  |
| ***R. envalirensis*** | <0.001 | <0.001 | 0.90 |  |  |  |  |  |  |  |
| ***R. marsicus*** | 0.11 | <0.01 | 1.00 | 0.89 |  |  |  |  |  |  |
| ***R. mediocompositus*** | <0.01 | <0.001 | 1.00 | 1.00 | 1.00 |  |  |  |  |  |
| ***R. austroslovenicus*** | <0.01 | <0.001 | 1.00 | 1.00 | 1.00 | 1.00 |  |  |  |  |
| ***R. subcarniolicus*** | 0.05 | <0.001 | 1.00 | 1.00 | 1.00 | 1.00 | 1.00 |  |  |  |
| ***R. notabilis*** | 1.00 | 0.90 | <0.01 | <0.001 | 0.06 | <0.001 | <0.001 | 0.02 |  |  |
| ***R. peracris*** | <0.01 | <0.001 | 0.42 | 1.00 | 0.47 | 1.0 | 1.00 | 1.00 | <0.01 |  |
| **mean** | 8.14 | 12.67 | 0.22 | 1.01 | 0.00 | 0.67 | 1.22 | 0.71 | 9.50 | 1.27 |

**B**

|  | ***R. cassubicifolius* s.l.** | ***R. flabellifolius*** | ***R. envalirensis* s.l.** | ***R. marsicus*** | ***R. notabilis* s.l.** |
| --- | --- | --- | --- | --- | --- |
| ***R. flabellifolius*** | <0.001 |  |  |  |  |
| ***R. envalirensis* s.l.** | <0.001 | 0.17 |  |  |  |
| ***R. marsicus*** | <0.01 | 0.55 | 0.17 |  |  |
| ***R. notabilis* s.l.** | <0.001 | 0.10 | 0.17 | 0.14 |  |
| **mean** | 12.12 | 0.22 | 1.01 | 0.00 | 2.25 |
