## Supplementary Text 1 for "Phylogenomics supported by geometric morphometrics reveals delimitation of sexual species within the polyploid apomictic *Ranunculus auricomus* complex (Ranunculaceae)"

**a) Library preparation and Hybrid Capture protocol for Target Enrichment.---**Genomic DNA was extracted from ~ 1.5 cm² leaf material of silica-dried samples or herbarium collection. For the scope, we used the Qiagen DNeasy Plant Mini Kit® (Qiagen, Hilden, Germany) following the manufacturer’s instructions, except for the sample incubation time in the lysis buffer that was increased to one hour. DNA quality and fragments length were checked by gel electrophoresis in a 1.5 Agarose gel and using the Roti®-Load DNAstain 3 (Carl Roth, Karlsruhe, Germany), particularly for herbarium specimens. Extract concentration was estimated using the Qubit® fluorometer and the Qubit® dsDNA HS Assay Kit (ThermoFisher Scientific, Waltham, USA). Sequencing libraries were prepared using either the ‘NEBNext Ultra II DNA Library Prep Kit for Illumina®’ (E7645) or the ‘NEBNext Ultra II FS DNA Library Prep Kit for Illumina®’ (E7805) (New England BioLabs, Ipswich, USA). In the former case, we shared DNA with a Bioruptor® Pico (Diagenode, Seraing, Belgium) before library preparation. Extracts were diluted to 10 ng/µL and sonicated for eight cycles of 15’ sonication and 90” break in order to obtain fragments of approximately 300-500 bp. In the latter case, enzymatic shearing is combined with the first steps of the library preparation. Fragmentation was carried out for 12’ at 37°C in order to obtain DNA fragments of the same length of the sonicated samples. For the herbarium collections Hoe5615 and Du33351-15, sharing incubation was shorter, 3’ and 10’, respectively. In both cases, we followed the manufacturer’s instructions. At the end of the library preparation procedure, samples were PCR amplified for 14 cycles during which sample-specific dual indices (‘NEBNext Multiplex Oligos for Illumina®’, E7600; New England BioLabs, Ipswich, USA) were added to the fragments. Indexed samples were pooled in equal quantities (four samples per 500 ng pool), dehydrated in a Concentrator Plus (Eppendorf, Hamburg, Germany) and diluted in 7 µL of ddH_2_O. Each pool was enriched using the custom baits kit following the manufacturer’s protocol. Hybridization took place for 21 h at 65 °C.

Enriched products were PCR-amplified for 14 cycles using the 2X KAPA HiFi HotStart Mix (KAPA Biosystems, Wilmington, USA) and the P7 and P5 adapters as primers. Amplified enriched libraries were purified with 50 µL of AMPure XP Beads (New England BioLabs, Ipswich, USA) following the manufacturer’s protocol. Concentrations were measured with the Qubit® fluorometer and fragment length distributions were checked with a Bioanalyzer (Agilent, Santa Clara, USA). In the few cases in which fragment length was different from the desired one (and especially in cases short fragments were present), pools were undergone to side selection with the BluePippin (Sage Science, Beverly, USA).

Sequencing took place on an Illumina MiSeq System (Illumina Inc., San Diego, USA) at the Transcriptome and Genome Analysis Laboratory (Georg-August-Universität, Göttingen, Germany). Pools were mixed equimolarly and sequenced in two different paired-end runs (6 pools, 24 samples each) with a 2 x 250 bp (500 cycles) v2 kit.

**b) Read processing und alignments.---**We checked the quality of raw reads with FastQC (available at: http://www.bioinformatics.bbsrc.ac.uk/projects/fastqc). Further processing of the raw reads was done using the pipeline HybPhyloMaker (all scripts available at: https://github.com/tomas-fer/HybPhyloMaker/; Fér & Schmickl, 2018). This pipeline offers a set of bash scripts assisting the procedure of assembly of captured sequences from read quality-trimming to the reconstruction of phylogenetic trees. In HybPhyloMaker, quality-trimmed individual raw reads are mapped to a reference sequence and then merged into contigs that are aligned for each gene separately. As pseudo-reference for read mapping, we used a sequence consisting of the concatenation of the target exonic sequences separated by stretches of 800 Ns. Sequence adapters were removed and reads were quality-trimmed using Trimmomatic vers. 0.32 (Bolger & al., 2014) with the default settings used in HybPhyloMaker. Duplicated reads were removed with FastUniq vers. 1.1 (Xu & al., 2012). Mapping to the pseudo-reference genome was done with BWA (Li & Durbin, 2010). In order to avoid the loss of allelic information during the process of allele mapping and consensus sequence production, we took out of the HybPhyloMaker pipeline the *.bam files produced after mapping and phased them with SAMtools v0.1.19 (Li & al., 2009) using the commands ‘samtools calmd’ and ‘samtools phase’. The respective *.bai files were also duplicated and named consequently. The phased *.bam and *.bai files were then placed back in a the HybPhyloMaker working directory for further processing within the pipeline workflow. A new sample file with duplicate sample names was placed in the’/10rawreads/’. The pipeline was therefore resumed for the computing of the consensus sequences (allele-wise consensus sequences) by calling the script ‘HybPhyloMaker2_readmapping.sh’ but specifying ‘mapping=no’ in the setting file. Consensus sequences were produced with ConsensusFixer (available at: https://github.com/cbg-ethz/ConsensusFixer) since this is the only of the approaches available in HybPhyloMaker able to call ambiguity DNA codes in case of multiple bases per site in the mapped reads. For ConsensusFixer, we used the following settings: minimum relative abundance of the alternative base (‘plurality’ in the setting file of HybPhyloMaker) of 0.2 and a minimum read coverage for ambiguity calling (‘mincov’) of 5.

Consensus sequences were matched to sequences of the target exons to produce PSLX files using BLAT (Kent, 2002). Afterward, they were combined to produce exon-wise matrices with ‘assembled_exons_to_fastas.py’ (Weitemier & al., 2014). These were aligned with MAFFT vers. 7.029 (Katoh, 2013) using the default program settings. HybPhyloMaker performs two consecutive steps to check the alignments for missing data and filter out those that exceed certain levels. First, sequences with more than a certain percentage of Ns in an alignment (‘MISSINGPERCENT’ in the settings file) are deleted. We set this option to 40. Secondly, alignments with less than a certain percent of sequences (‘SPECIESPRESENCE’ in the settings file) are filtered out. We set this value to 75 so that alignments with more than 25% of missing sequences were excluded.
